# Linear ubiquitination at damaged lysosomes induces local NF-κB activation and controls cell survival

**DOI:** 10.1101/2023.10.06.560832

**Authors:** Laura Zein, Marvin Dietrich, Denise Balta, Verian Bader, Christoph Scheuer, Suzanne Zellner, Nadine Weinelt, Julia Vandrey, Muriel C. Mari, Christian Behrends, Friederike Zunke, Konstanze F. Winklhofer, Sjoerd J. L. van Wijk

## Abstract

Lysosomes are the major cellular organelles responsible for nutrient recycling and degradation of cellular material. Maintenance of lysosomal integrity is essential for cellular homeostasis and lysosomal membrane permeabilization (LMP), induced by lysosomotrophic agents, sensitizes towards cell death. Damaged lysosomes are repaired or degraded via lysophagy, during which glycans, exposed on ruptured lysosomal membranes, are recognized by galectins leading to K48- and K63-linked poly-ubiquitination (poly-Ub) of lysosomal proteins followed by recruitment of the autophagic machinery and degradation. Linear (M1) poly-Ub, catalyzed by the E3 ligase linear ubiquitin chain assembly complex (LUBAC) and removed by the OTU domain-containing deubiquitinase with linear linkage specificity (OTULIN) exerts important functions in immune signaling and cell survival, but the role of M1 poly-Ub in lysosomal homeostasis remains largely unexplored. Here, we demonstrate that damaged lysosomes are decorated with M1 poly-Ub in a LUBAC-, OTULIN- and K63-dependent manner. LMP-induced M1 poly-Ub at damaged lysosomes contributes to lysosome degradation, recruits nuclear factor κ-B (NF-κB) essential modulator (NEMO) and locally activates inhibitor of NF-ĸB kinase (IKK) to trigger NF-κB activation in a K63 poly-Ub-dependent manner. Inhibition of lysosomal degradation enhances LMP- and OTULIN-dependent cell death, indicating pro-survival functions of LMP and potentially lysophagy. Finally, we demonstrate that M1 poly-Ub occurs at L-leucyl-leucine methyl ester (LLOMe)-damaged lysosomes in primary mouse neurons and induced pluripotent stem cell (iPSC)-derived primary human dopaminergic neurons. Together, our results reveal novel functions of M1 poly-Ub during lysosomal homeostasis, LMP and degradation of damaged lysosomes, with important implications for NF-κB signaling, inflammation and cell death.

## Introduction

Lysosomes are acidic cellular organelles required for recycling essential nutrients through degradation of autophagic cargo, including proteins, nucleic acids, polysaccharides and lipids, endo- and phagocytosed materials, as well as damaged organelles, protein aggregates and intracellular pathogens [1]. In addition, lysosomes mediate plasma membrane repair, inflammation, lipid homeostasis and programmed cell death via fusion with endocytic vesicles or the release of lysosomal hydrolases, like cathepsin B and D. Damage to lysosomes induced by reactive oxygen species (ROS), sphingosine, lysosomotrophic agents, such as L-leucyl-leucine methyl ester (LLOMe) and cationic amphiphilic drugs (CADs), like amiodarone (AMIO), pimozide (PIMO) and loperamide (LOP), triggers lysosomal membrane permeabilization (LMP) and cytosolic release of cathepsins [2–5]. Lysosomal damage and LMP underlie lysosomal cell death (LCD), caused by the release and hydrolytic action of lysosomal enzymes in the cytosol [6]. Secondary LMP, cytosolic cathepsin release and lysosome-dependent cell death are also observed in final stages of other forms of cell death, including apoptosis and necroptosis [6].

Damaged lysosomes can either be repaired or degraded via lysophagy [7]. Lysophagy is typically initiated upon lysosomal damage and permeabilization, during which exposed glycosylated lysosomal transmembrane proteins (glycans) induce the recruitment of galectin-1 (LGALS1), −3 (LGALS3), −8 (LGALS8) and −9 (LGALS9) [1]. In addition, the E3 ligase tripartite motif 16 (TRIM16), the SCF E3 ligase complex subunit F-box only protein 27 (FBXO27) and the Ub-conjugating enzyme E2Q-like protein 1 (UBE2QL1) mediate K48- and K63-linked poly-Ub modification of lysosomal proteins in a p97/valosin-containing protein (VCP)- and the ubiquitin thioesterase OTU1 (YOD1)-dependent manner [8–11]. Poly-Ub chains subsequently recruit autophagy receptor proteins, like p62/Sequestosome-1 (SQSTM1), nuclear dot protein 52 (NDP52) and Tax1-binding protein 1 (TAX1BP1), that serve as connective linkers with membrane-anchored, lipidated autophagy-related protein 8 (ATG8) family members [12,13] to incorporate ubiquitinated lysosomes in microtubule-associated protein 1A/1B light chain 3 (MAP1LC3/LC3)-decorated phagophores [1,9].

Different types of poly-Ub chain linkages, including linear, K48 and K63, are implicated in selective autophagy of damaged organelles, intracellular pathogens and protein aggregates [14,15]. Linear, or M1, poly-Ub is generated by the linear ubiquitin chain assembly complex (LUBAC) E3 ligase and cleaved by the OTU domain-containing deubiquitinase (DUB) with linear linkage specificity (OTULIN). LUBAC is composed of Shank-associated RH domain-interacting protein (Sharpin), heme-oxidized IRP2 ubiquitin ligase 1 (HOIL1) and HOIL-1-interacting protein (HOIP/RNF31) [16–20]. As the catalytic subunit, HOIP is responsible for M1 poly-Ub via the RING1-IBR-RING2 (RBR) domain, containing the RBR E3 ligase-specific catalytic Cys885. Apart from attracting autophagy receptor proteins, M1 poly-Ub recruit NEMO and activate IKKα/β [21,22]. NEMO and IKK activation control the proteasomal stability of NF-ĸB inhibitor α (IĸBα) and mediate the release of NF-ĸB transcription factors to the nucleus to activate pro-survival gene expression programs [23,24]. Tank-binding kinase 1 (TBK1) is also activated at ubiquitinated cargo and further activates NF-κB. At the same time, activated TBK1 phosphorylates autophagy receptors such as p62/SQSTM1, optineurin (OPTN) or NDP52 [25–27] and promotes receptor interactions with poly-Ub and ATG8 proteins, recruitment of the autophagy machinery and phagophore formation [28–30]. By doing so, TBK1 contributes to different forms of selective autophagy, including mitophagy, xenophagy and lysophagy [25,26,29,31,32].

Even though the role of M1 poly-Ub has already been characterized in the autophagic clearance of bacteria (xenophagy) [33], the functions of M1 poly-Ub in other forms of selective autophagy, such as lysophagy, remain unclear. Here, we demonstrate that during LMP, lysosomes become modified with M1 poly-Ub in a LUBAC- and OTULIN-dependent manner, contributing to the degradation of damaged lysosomes. M1 poly- Ub-modified lysosomes initiate NF-κB signaling by recruiting NEMO and local activation of IKK. Mechanistically, we demonstrate that deposition of lysosomal M1 poly-Ub relies on pre-formed K63-linked poly-Ub and loss of K63 poly-Ub at damaged lysosomes prevents M1 poly-Ub and NF-κB activation. Interfering with autophagic degradation by cathepsin inhibition or via loss of the major autophagy receptor proteins increases LMP-induced cell death, indicating that lysophagy serves pro-survival functions in response to lysosomal damage. Finally, we demonstrate modification of damaged lysosomes by M1 poly-Ub in primary murine cortical neurons and in differentiated induced pluripotent stem cell (iPSC)-derived human dopaminergic neurons. Together, our findings identify new and emerging functions of M1 poly-Ub at damaged lysosomes with important implications for lysosome homeostasis, inflammation and cell survival in health and disease.

## Results

### M1 poly-Ub regulates the autophagic flux and sequestration of specific autophagic cargo

Both LUBAC and OTULIN have been associated with the regulation of the autophagic flux and cargo selection during selective autophagy [33,34], but the question as to how M1 poly-Ub provides cargo selectivity remains unresolved. To characterize M1 poly-Ub-specific cargo during (selective) autophagy, loss of OTULIN was established in glioblastoma (GBM) MZ-54 cells using CRISPR/Cas9-mediated knockout (KO). Of note, GBM cells display high levels of basal autophagy activity [35,36] and serve as ideal models to study (selective) autophagy pathways [37]. Loss of OTULIN increased basal M1 poly-Ub levels, accompanied with a reduction of the levels of the LUBAC subunits HOIP and HOIL1, but not Sharpin **(Figure 1A)**. In addition, increased M1 poly- Ub upon loss of OTULIN slightly increased basal LC3B lipidation that was further amplified in response to autophagy activation with the mammalian target of rapamycin (mTOR) complex 1 (mTORC1) inhibitor Torin-1, or with the autophagy-inducing substance loperamide (LOP) [38] **(Figure 1B and 1C)**. This was accompanied by OTULIN-dependent increases in LC3- and M1 poly-Ub-positive puncta upon treatment with LOP **(Figure 1D and 1E)**. Loss of OTULIN slightly increased the basal punctate accumulation of M1 poly-Ub, that potently increased upon LOP treatment **(Figure 1E)**. Blockage of lysosomal acidification and subsequent degradation of autophagic cargo with the vacuolar H^+^-ATPase inhibitor bafilomycin A1 (BafA1) increased LC3-II accumulation in OTULIN KO cells, compared to controls **(Figure 1F)**, indicating OTULIN-dependent increases in autophagy flux. To investigate the role of OTULIN in autophagy in more detail, control and OTULIN KO cells were subjected to electron microscopy analysis. Intriguingly, OTULIN-deficient cells displayed a striking increase in basal and LOP-induced degradative compartments comprising lysosomes, amphisomes and autolysosomes and, to a lesser extent, also in autophagosomes compared to control cells **(Figure 1G and 1H)**. To identify OTULIN-dependent cargo, myc-APEX2-LC3B-expressing control and OTULIN-deficient cells were generated **(Supplemental Figure 1A)**, incubated with biotin-phenol and pulsed with H_2_O_2_ to allow biotinylation of proteins in close proximity of LC3B. This was followed by proteinase K-mediated proteolysis of non-enclosed proteins, streptavidin pulldown and mass spectrometry [39]. Proteinase K digestion in the absence and presence of the membrane-lysing agent Triton-X-100 validated autophagosomal protection of biotinylated proteins, myc-APEX2-LC3B and endogenous p62/SQSTM1 **(Supplemental Figure 1B)**. In contrast, the ATG8-interacting protein Kelch repeat and BTB domain-containing protein 7 (KBTBD7) was degraded in the presence of proteinase K **(Supplemental Figure 1B)**, as described earlier [40]. In addition, LOP decreased the levels of biotinylated proteins, reflecting autophagic degradation of biotinylated proteins, whereas autophagy blockade with BafA1 increased the levels of biotinylated proteins **(Supplemental Figure 1C)**. Mass spectrometry of biotinylated fractions from BafA1-treated control and OTULIN KO cells revealed prominent alterations in autophagosome content and autophagic cargo **(Figure 1I)**, including several organelle-specific proteins (e.g. VDAC2, ERLIN1, TOMM40), endosomal and lysosomal proteins (e.g. RAB7A, RAB11A), TAX1BP1 and LGALS3. Validation with proteinase K protection assays verified selected hits as autophagosomal membrane cargo **(Supplemental Figure 1D)**. The autophagy receptor TAX1BP1 is implicated in many forms of selective autophagy, including xenophagy, aggrephagy, mitophagy and lysophagy [31,41–43] and autophagy receptors are co-degraded with their cargo. To validate OTULIN-dependent TAX1BP1 degradation, we tested the effect of BafA1 treatment on TAX1BP1 protein abundance. Indeed, BafA1 increased TAX1BP1 levels in OTULIN KO cells compared to controls **(Figure 1J)**. Enhanced BafA1-mediated accumulation of TAX1BP1 in OTULIN KO cells suggests OTULIN-dependent differences in the autophagic degradation of TAX1BP1. TAX1BP1 was degraded faster in cycloheximide (CHX)-treated OTULIN KO cells compared to controls and this degradation could be blocked by BafA1, indicating OTULIN-dependent turnover of TAX1BP1 via autophagy **(Supplemental Figure 1E and 1F)**. Taken together, these data reveal that OTULIN negatively regulates the autophagic flux and controls the degradation of TAX1BP1 and potentially TAX1BP1-associated autophagic cargo.

**Figure 1.**
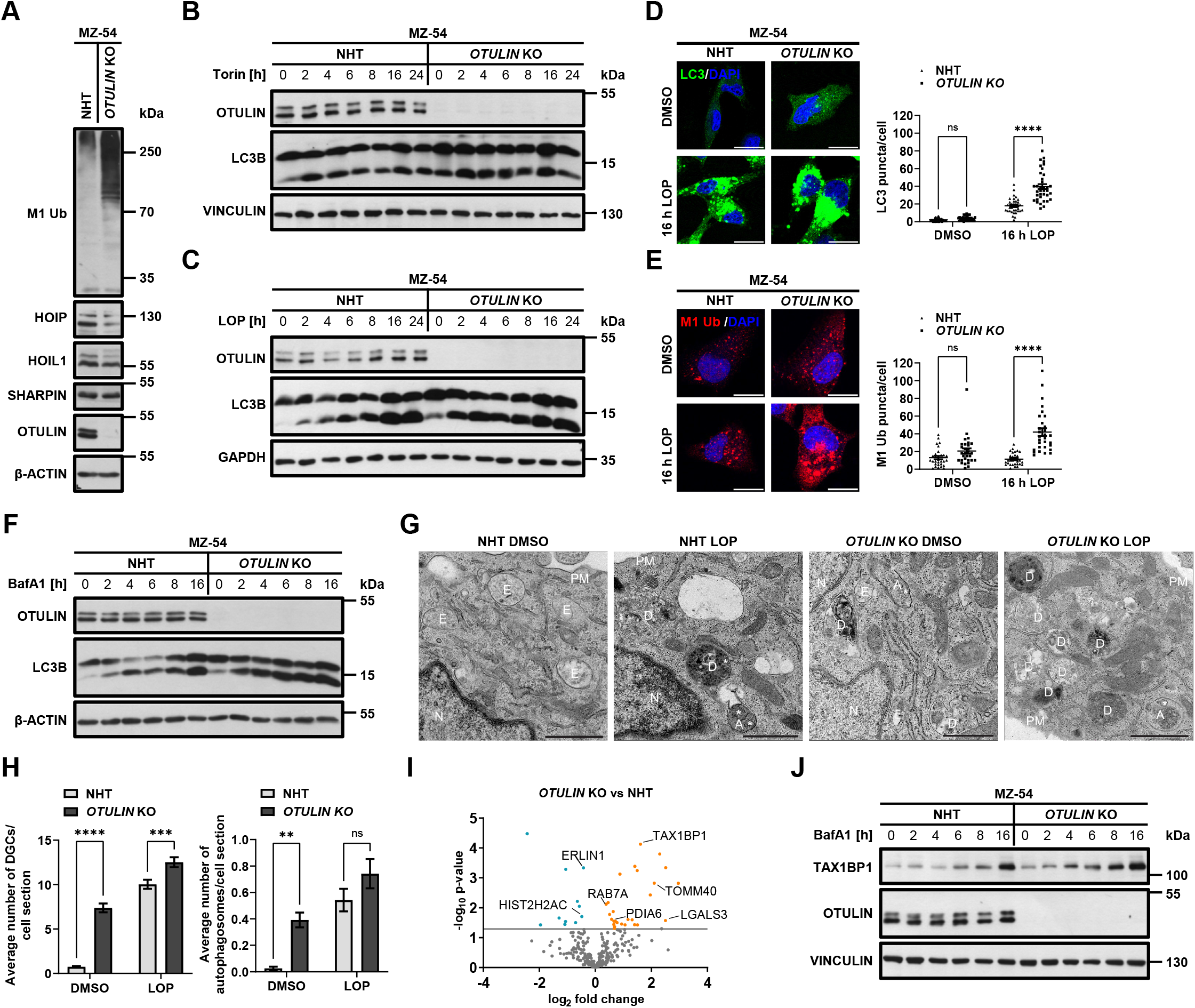
OTULIN negatively regulates the autophagic flux and sequestration of specific autophagic cargo. A. Western blot analysis of total M1 poly-Ub, HOIP (∼119 kDa), HOIL1 (∼57 kDa), Sharpin (∼40 kDa) and OTULIN levels in MZ-54 non-human target (NHT) and OTULIN KO GBM cells. β-actin was used as loading control. Representative blots of at least two independent experiments are shown. **B.** Western blot analysis of LC3B lipidation in MZ-54 NHT and OTULIN KO GBM cells, treated with Torin-1 (0.25 µM) for the indicated time points. Vinculin was used as loading control. Representative blots of at least two independent experiments are shown. **C.** Idem as B, but cells were treated with 17.5 µM loperamide (LOP). GAPDH was used as loading control. Representative blots of at least two independent experiments are shown. **D.** Representative immunofluorescence staining of LC3B in MZ-54 NHT and OTULIN KO GBM cells, treated with LOP (17.5 µM) for 16 h. Scale bar: 15 µm. Right panel: Quantification of LC3B puncta. Mean and SEM of 36 cells from three independent experiments are shown. **** p < 0.0001, ns: not significant. **E.** Idem as D, but MZ-54 NHT and OTULIN KO GBM cells treated with LOP (17.5 µM) for 16 h were stained for M1 poly-Ub. Scale bar: 15 µm. Right panel: Quantification of M1 Ub puncta. Mean and SEM of 30 cells from three independent experiments are shown. **** p < 0.0001, ns: not significant. **F.** Idem as B, but cells were treated with 200 nM BafA1 for the indicated time points. β-actin was used as loading control. Representative blots of at least two independent experiments are shown. **G.** Representative electron microscopy images of MZ-54 NHT and OTULIN KO cells treated with LOP (17.5 µM) for 16 h. A, autophagosome; D, degradative compartment; ER, endoplasmic reticulum; M, mitochondria; N, nucleus; PM, plasma membrane; asterisk, rough-ER fragment. Scale bar: 1 µm. **H.** Quantification of the average number of DGCs and autophagosomes per cell section of G, described in the Methods section. Mean and SEM are shown. ** p < 0.01, *** p < 0.001, **** p < 0.0001, ns: not significant. **I.** Volcano plot of significantly up- and downregulated (orange and blue, respectively) autophagosome enriched cargo candidates from myc-APEX2-LC3B-expressing MZ-54 NHT and OTULIN KO cells treated with BafA1 (200 nM) for 2 h, followed by biotinylation, proteinase K digestion, streptavidin pulldown and mass spectrometry. Data are derived from four replicates. p ≤ 0.05. **J.** Western blot analysis of TAX1BP1 and OTULIN expression in MZ-54 NHT and OTULIN KO GBM cells, treated with BafA1 (200 nM) for the indicated time points. Vinculin was used as loading control. Representative blots of at least two independent experiments are shown. Statistical significance was determined with two- way analysis of variance (ANOVA) followed by Tukey’s multiple comparisons tests (**D&E**), and with student’s t-test (**I**).

### OTULIN regulates M1 poly-Ub accumulation at damaged lysosomes

TAX1BP1 serves as autophagy receptor for the degradation of damaged lysosomes. Besides OTULIN-dependent enrichment of TAX1BP1 in autophagosomes, LGALS3 was also enriched, raising the possibility that OTULIN and M1 poly-Ub might affect lysosomal homeostasis. To address the role of OTULIN in lysophagy, LMP and lysosomal degradation was induced with the lysosomotrophic drug LLOMe. Surprisingly, LLOMe strongly induced global M1 poly-Ub levels in OTULIN-deficient cells, accompanied by LLOMe- and OTULIN-dependent alterations in degradation of LGALS3 and LC3 lipidation **(Figure 2A)**. Immunofluorescence analysis confirmed OTULIN- and LLOMe-induced M1 poly-Ub accumulation at damaged lysosomes **(Figure 2B and 2C)**.

**Figure 2.**
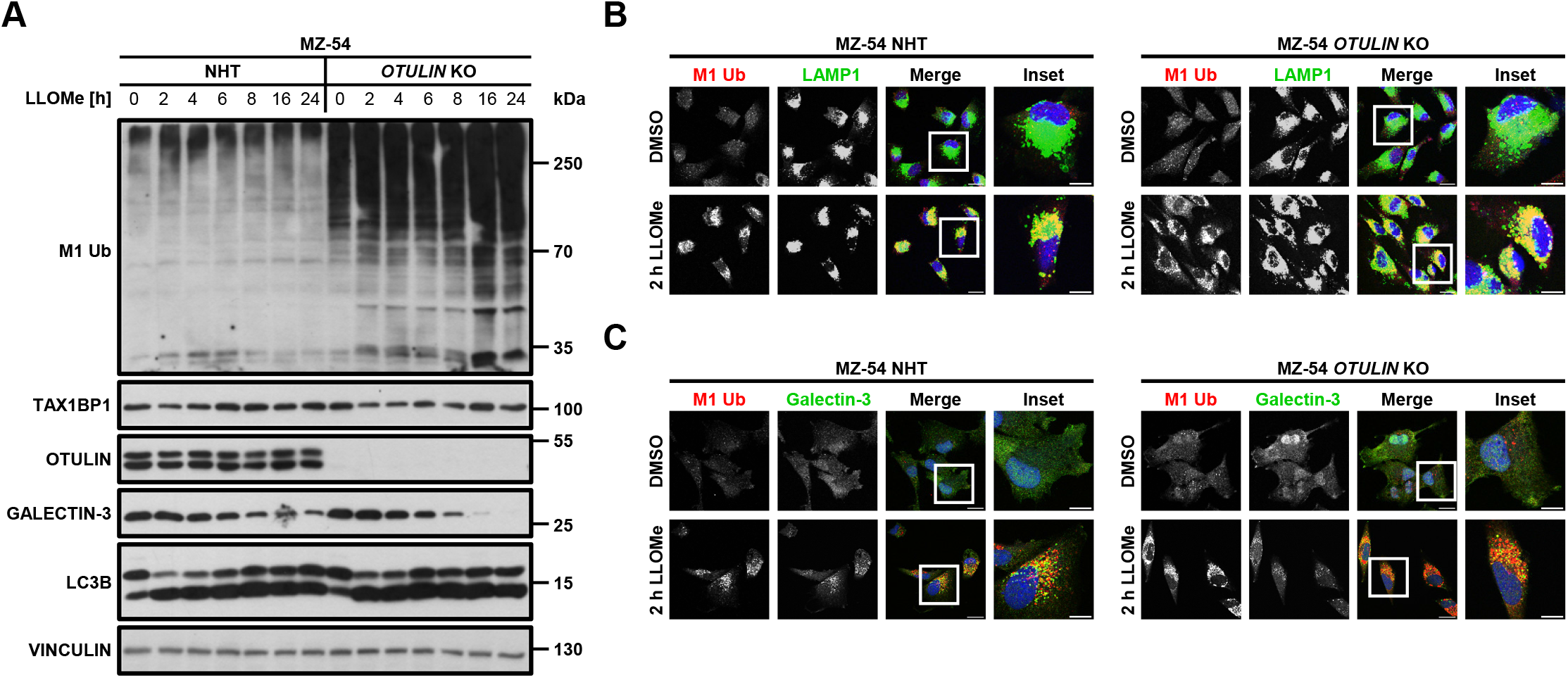
M1 poly-Ub accumulates at lysosomes in response to LLOMe-induced damage. A. Western blot analysis of M1 poly-Ub, TAX1BP1, OTULIN, LGALS3 and LC3B expression in MZ-54 NHT and OTULIN KO GBM cells, treated with LLOMe (500 µM) for the indicated time points. Vinculin was used as loading control. Representative blots of at least two independent experiments are shown. **B.** Representative immunofluorescence staining of M1 poly-Ub and LAMP1 in MZ-54 NHT and OTULIN KO GBM cells, treated with LLOMe (500 µM) for 2 h. Scale bar: 20 µm (10 µm for insets). **C.** Idem as B, but MZ-54 NHT and OTULIN KO GBM cells treated with LLOMe (500 µM) for 2 h were stained for M1 poly-Ub and LGALS3. Scale bar: 20 µm (10 µm for insets).

In agreement with the autophagosome content profiling, TAX1BP1 was also recruited to LLOMe-damaged lysosomes and partially colocalized with lysosome-associated membrane protein 1 (LAMP1) and M1 poly-Ub **(Supplemental Figure 2A)**. To investigate if the autophagic clearance of damaged lysosomes relies on OTULIN and M1 poly-Ub, immunofluorescence of LGALS3 and LAMP1 was performed after LLOMe treatment and washout **(Supplemental Figure 2B)**. Intriguingly, degradation of LGALS3 puncta was only slightly enhanced in OTULIN-deficient cells compared to controls, indicating that OTULIN and M1 poly-Ub might have a minor impact on lysophagy **(Supplemental Figure 2C)**. Since TBK1 is activated in response to lysosomal damage and is required for the lysophagic flux [26,31], we investigated TBK1 activation upon LLOMe-induced lysosomal damage. Interestingly, TBK1 phosphorylation at S172 was increased in LLOMe-treated OTULIN-deficient cells compared to controls **(Supplemental Figure 2D)** and this was partially blocked by HOIPIN-8-mediated inhibition of HOIP [44] **(Supplemental Figure 2E)**, suggesting M1 poly-Ub-dependent regulation of TBK1 activation at damaged lysosomes. To investigate the broader relevance of M1 poly-ubiquitination in the cell-autonomous response to lysosomal damage, we performed LLOMe treatment in HeLa cells and this also triggered formation of LGALS3 puncta and accumulation of M1 poly-Ub at damaged lysosomes **(Supplemental Figure 2F and 2G)**. Together, these results demonstrate that lysosomes are marked with M1 poly-Ub in response to LLOMe-induced damage in an OTULIN-dependent manner.

### OTULIN-mediated accumulation of M1 poly-Ub at damaged lysosomes locally activates NF-κB

LUBAC- and OTULIN-regulated M1 poly-Ub controls xenophagy and accumulation of M1 poly-Ub at cytosolic bacteria was shown to create local NF-κB signaling platforms [33]. Recently, NEMO has been shown to be recruited to damaged mitochondria in a Parkin-dependent manner to activate NF-κB signaling, although M1 poly-Ub did not play functional roles in NEMO recruitment [45]. To investigate if M1 poly-Ub-modified lysosomes serve as platforms for local NF-κB activation, we performed immunofluorescence and confirmed recruitment of NEMO and activation of IKK at LLOMe-induced M1 poly-Ub-decorated lysosomes in MZ-54 and HeLa cells **(Figure 3A, 3B, 3C, Supplemental Figure 3A and 3B)**. Inhibition of HOIP activation with HOIPIN-8 reduced the punctate NEMO accumulation at damaged and M1 poly-Ub-modified lysosomes **(Figure 3D and Supplemental Figure 3C)**. Similarly, inhibition of IKKβ with TPCA-1 [46] also blocked the local activation of IKK at M1 Ub-modified damaged lysosomes **(Figure 3E and Supplemental Figure 3D)**. Intriguingly, local activation of NF-κB at M1-poly-Ub-modified damaged lysosomes strongly induced the expression of NF-κB target genes *IL6*, *IL8* and *TNFA* in an OTULIN-dependent manner, but not in control cells **(Figure 3F)**. OTULIN- and LLOMe-induced NF-κB target gene transcription was completely reduced by incubating with HOIPIN-8 or TPCA-1 **(Figure 3F)**, suggesting central roles for LUBAC and IKK in OTULIN- and M1 poly-Ub-dependent LMP-induced NF-κB activation.

**Figure 3.**
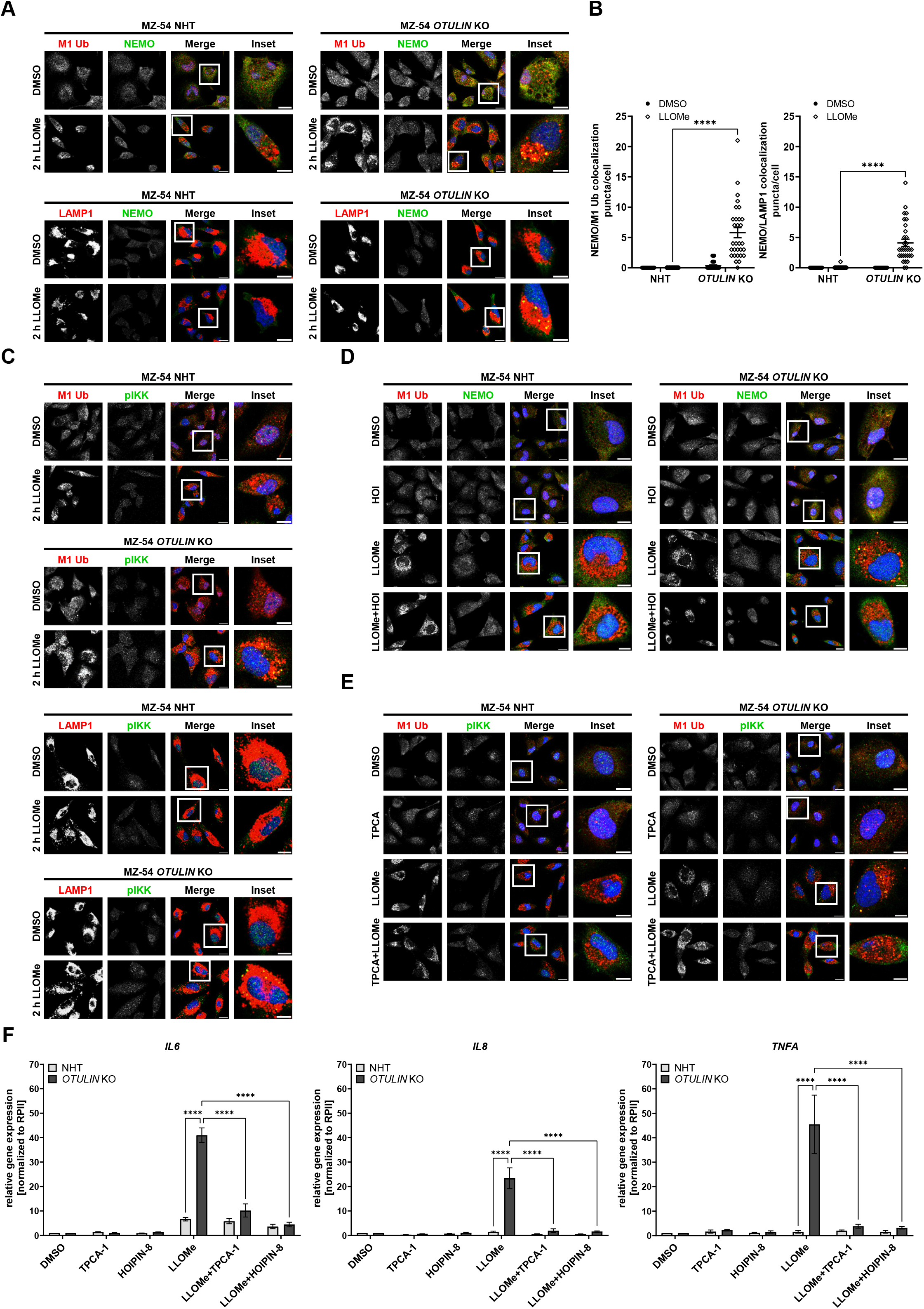
Lysosomal accumulation of M1 poly-Ub locally induces NF-κB activation during lysophagy. A. Representative immunofluorescence staining of M1 poly-Ub/NEMO (upper panel) and LAMP1/NEMO (lower panel) in MZ-54 NHT and OTULIN KO GBM cells, treated with LLOMe (500 µM) for 2 h. Scale bar: 20 µm (10 µm for insets). **B.** Quantification of M1 poly-Ub- and NEMO-positive and M1 poly Ub- and LAMP1-positive puncta from A. Mean and SEM of 30-31 cells from four independent microscopic views are shown. **** p < 0.0001. **C.** Idem as A, but MZ-54 NHT and OTULIN KO GBM cells treated with LLOMe (500 µM) for 2 h were stained for M1 poly-Ub/pIKKα/β (Ser176/180) and LAMP1/pIKKα/β (Ser176/180). Scale bar: 20 µm (10 µm for insets). **D.** Idem as A, and C but MZ-54 NHT and OTULIN KO GBM cells pre-treated with HOIPIN-8 (30 µM) for 1 h before addition of LLOMe (500 µM) for 2 h were stained for M1 poly-Ub and NEMO. Scale bar: 20 µm (10 µm for insets). **E.** Idem as A, C and D, but MZ-54 NHT and OTULIN KO GBM cells pre-treated with TPCA-1 (5 µM) for 1 h before addition of LLOMe (500 µM) for 2 h were stained for M1 poly-Ub and pIKKα/β (Ser176/180). Scale bar: 20 µm (10 µm for insets). **F.** mRNA expression levels of *IL6*, *IL8* and *TNFA* were analyzed by qRT-PCR of MZ-54 NHT and OTULIN KO GBM cells, pre-treated with HOIPIN-8 (30 µM) and TPCA-1 (5 µM) for 1 h and treated with LLOMe (500 µM) for 4 h. Fold increase of mRNA levels is shown relative to DMSO control with mean and SEM of at least three independent experiments performed in triplicates. **** p < 0.0001, ns: not significant. Statistical significance was determined with two-way analysis of variance (ANOVA) followed by Tukey’s multiple comparisons tests (**B&F**).

### Local formation of M1 Ub chains and NF-κB activation relies on K63 poly-Ub deposited on damaged lysosomes

In other cellular contexts, the recruitment of LUBAC to activated death receptors requires pre-modification with K63-linked poly-Ub [47,48]. To address the requirement of pre-deposited K63 poly-Ub for M1 poly-Ub modification of damaged lysosomes, the dimeric K63 poly-Ub-specific E2 enzyme UbcH13-UEV1a was inhibited with NSC697923 upon the induction of LLOMe-induced lysosomal damage and lysophagy. NSC697923 strongly reduced LLOMe-triggered increased K63-linked poly-Ub at damaged lysosomes **(Figure 4A)**, confirming inhibition of the K63 poly-Ub machinery. Intriguingly, inhibition of UbcH13-UEV1A also significantly reduced LLOMe-induced M1 poly-Ub at damaged lysosomes, suggesting that M1 poly-Ub modification of damaged lysosomes might critically rely on pre-deposited K63 poly-Ub **(Figure 4B)**. In addition, NSC697923 also potently inhibited accumulation of NEMO at damaged lysosomes in LLOMe-treated OTULIN KO cells **(Figure 4C and 4D)** and reduced OTULIN- and LLOMe-dependent upregulation of NF-κB target genes **(Figure 4E)**.

**Figure 4.**
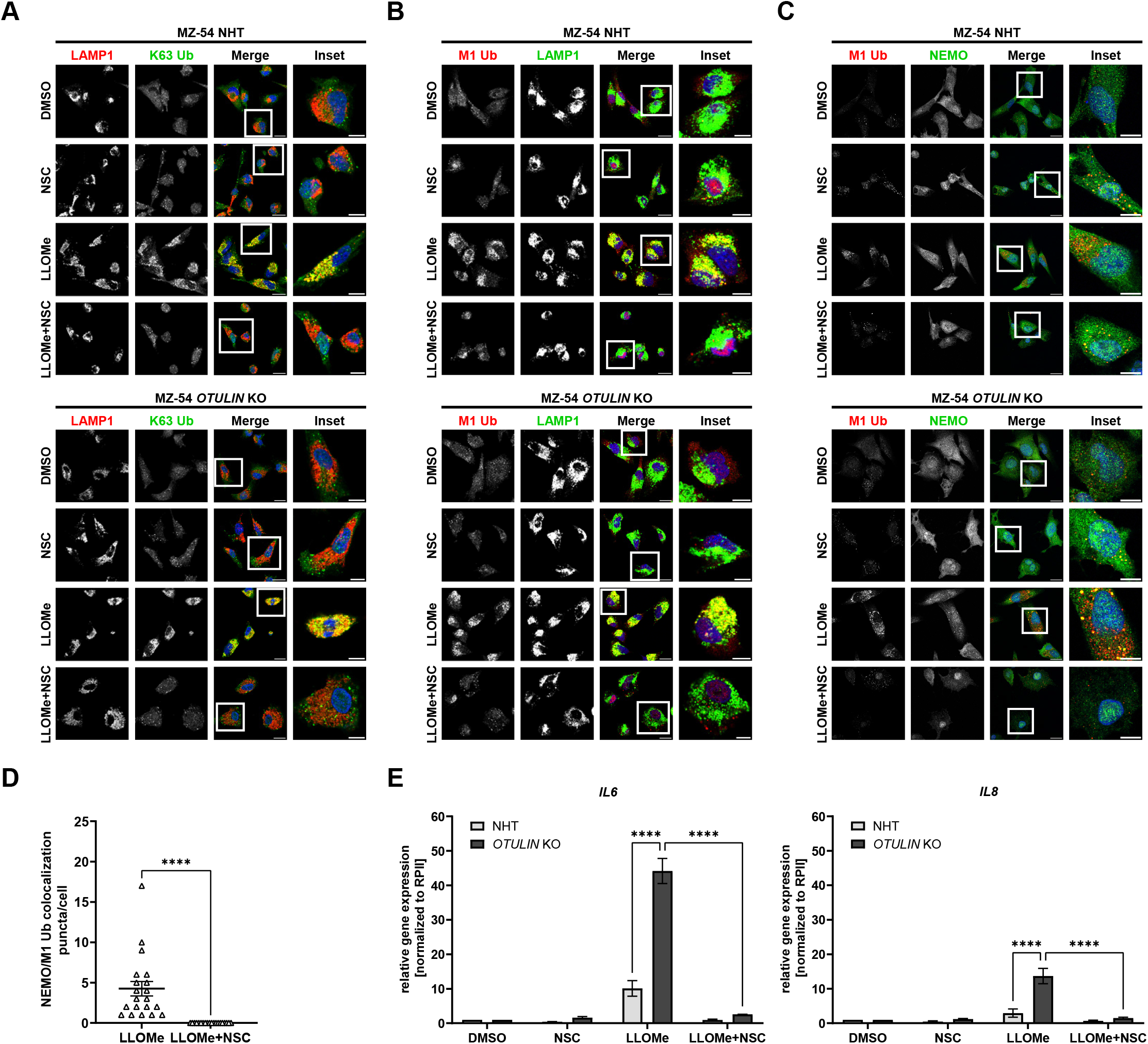
Formation of M1 poly-Ub chains and NF-κB activation at damaged lysosomes relies on K63 poly-Ub. A. Representative immunofluorescence staining of LAMP1 and K63 poly-Ub in MZ-54 NHT (upper panel) and OTULIN KO (lower panel) GBM cells pre-treated with NSC697923 (2.5 µM) for 1 h and treated with LLOMe (500 µM) for 2 h. Scale bar: 20 µm (10 µm for insets). **B.** Idem as A but MZ-54 NHT and OTULIN KO GBM cells were stained for M1 poly-Ub and LAMP1. Scale bar: 20 µm (10 µm for insets). **C.** Idem as A and B but MZ-54 NHT and OTULIN KO GBM cells were stained for M1 poly-Ub and NEMO. Scale bar: 20 µm (10 µm for insets). **D.** Quantification of M1 poly-Ub- and NEMO-positive puncta from C. Mean and SEM of 20 cells from four independent microscopic views are shown. **** p < 0.0001. **E.** mRNA expression levels of *IL6* and *IL8* were analyzed by qRT-PCR of MZ-54 NHT and OTULIN KO GBM cells pre-treated with NSC697923 (2.5 µM) for 1 h and treated with LLOMe (500 µM) for 4 h. Fold increase of mRNA levels is shown relative to DMSO control with mean and SEM of at least three independent experiments performed in triplicates. **** p < 0.0001, ns: not significant. Statistical significance was determined with unpaired t-test (**D**) and with two-way analysis of variance (ANOVA) followed by Tukey’s multiple comparisons tests (**E**).

### OTULIN-mediated M1 poly-Ub modification of damaged lysosomes promotes cell survival

The mass spectrometry-based analysis of OTULIN-dependent autophagic cargo revealed increased LGALS3 in autophagosomes, suggesting that OTULIN fulfills basal homeostatic functions at lysosomes. Since severe LMP might eventually activate cell death pathways, like apoptosis [49], we next sought to investigate LMP-induced cell death pathways. LLOMe treatment triggered cell death which could be blocked by caspase inhibition with zVAD.fmk in both control and OTULIN KO cells **(Figure 5A)**, confirming that LLOMe induces caspase-dependent apoptosis [50]. Since damaged lysosomes can either be repaired or degraded via lysophagy in a highly dynamic fashion to control the balance between cell survival and lysosome-dependent apoptotic cell death [9,51], increased M1 poly-Ub-mediated NF-kB activation at damaged lysosomes might fulfill cell death-regulating functions. If OTULIN-mediated M1 poly-Ub regulates lysosome-dependent apoptotic cell death, prevention of autophagic degradation of damaged lysosomes by either inhibition of cathepsin B and D or loss of autophagy receptor proteins, including those involved in lysophagy, should increase cell death. To test this idea, siRNA-mediated knock-down of OTULIN expression was performed in wild-type (WT) and Penta KO HeLa cells, deficient of the autophagy receptors OPTN, NDP52, TAX1BP1, p62/SQSTM-1 and Next to BRCA1 gene 1 protein (NBR1))[43], followed by inhibition of aspartyl proteases with Pepstatin A and the cathepsin B inhibitor CA-074 and LLOMe treatment **(Figure 5B and 5C)**. Already in the absence of LLOMe, Penta KO cells are more sensitive towards Pepstatin A/CA-074- and OTULIN-dependent cell death compared to siCtrl cells **(Figure 5B)**. Intriguingly, combined LLOMe-mediated lysosomal damage and cathepsin inhibition potently triggered cell death and was further enhanced after silencing of OTULIN expression in both WT and Penta KO cells **(Figure 5B)**, suggesting pro-survival roles of OTULIN-mediated M1 poly-Ub at damaged lysosomes.

**Figure 5.**
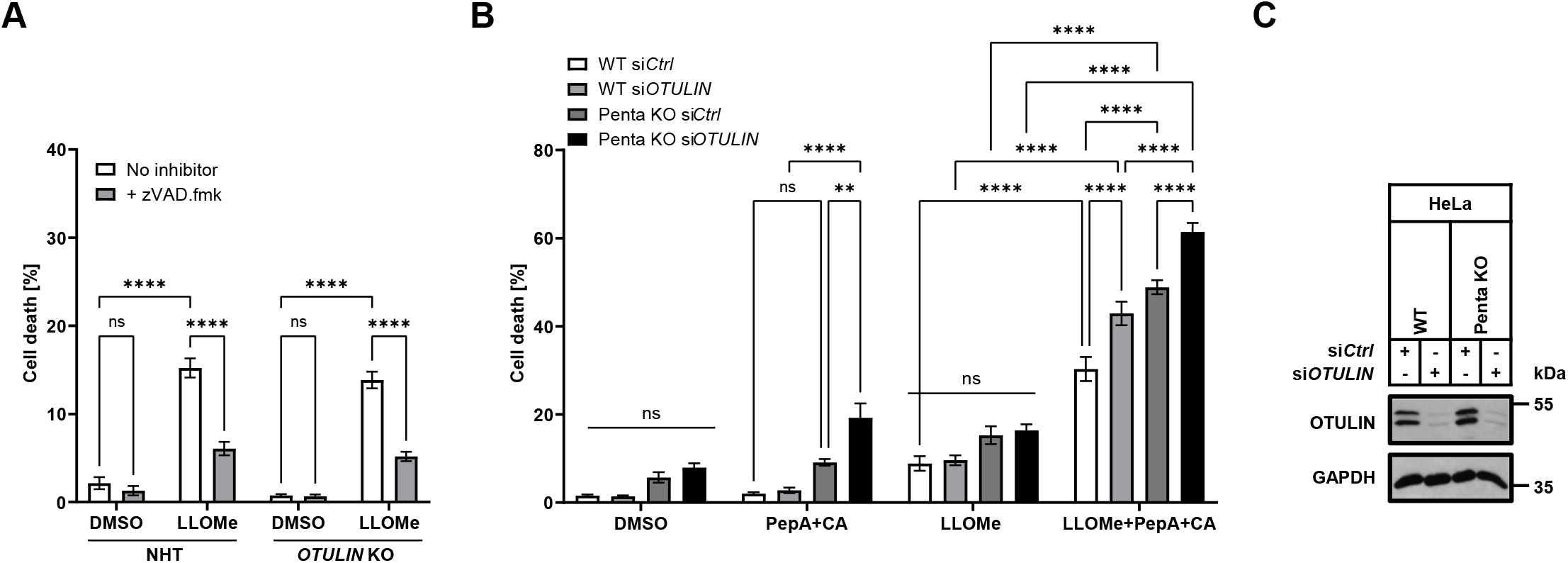
Cathepsin inhibition sensitizes GBM cells to LLOMe-induced apoptosis and lysophagy protects against cell death. A. Cell death analysis of MZ-54 NHT and OTULIN KO GBM cells treated with LLOMe (2 mM) and zVAD.fmk (20 µM) for 24 h. Cell death was assessed by measuring the propidium iodide (PI) uptake as fraction of total nuclei determined by Hoechst counterstaining using high-content fluorescence microscopy. Mean and SEM of three independent experiments are shown. **** p < 0.0001, ns: not significant. **B.** Cell death analysis after knockdown of OTULIN in HeLa WT and Penta KO cells and treatment with LLOMe (2 mM) and Pepstatin A (10 µM) and CA-074 (10 µM) for 24 h. Cell death was assessed by measuring the propidium iodide (PI) uptake as fraction of total nuclei determined by Hoechst counterstaining using high-content fluorescence microscopy. Mean and SEM of three independent experiments are shown. ** p < 0.01, **** p < 0.0001, ns: not significant. **C.** Western blot analysis of OTULIN expression in HeLa WT and Penta KO cells after knockdown of OTULIN to confirm silencing of protein expression. GAPDH was used as loading control. Representative blots of three independent experiments are shown. Statistical significance was determined with two-way analysis of variance (ANOVA) followed by Tukey’s multiple comparisons tests (**A&B**).

### M1 poly-Ub accumulates at damaged lysosomes in primary mouse neurons and human iPSC-derived dopaminergic neurons

Since lysosomal dysfunction is characteristic for many neurodegenerative diseases, including Parkinson’s disease, Alzheimer’s disease and amyotrophic lateral sclerosis (ALS) [52–54], we next investigated the role of M1 poly-Ub after lysosomal damage in human induced pluripotent stem cells (iPSC)-derived dopaminergic neurons (DA-iPSn). Successful differentiation into dopaminergic neurons was confirmed by Western blot analysis of the neuronal markers tyrosine hydroxylase (TH), neurofilament and Synapsin-I **(Supplemental Figure 4)**. Immunofluorescence analysis showed that LLOMe triggered the accumulation of M1 poly-Ub and localization of LGALS3 puncta at M1 poly-Ub modified damaged lysosomes in DA-iPSn **(Figure 6 A and B)**. Finally, primary mouse embryonic cortical neurons at 7 days *in vitro* (DIV) were incubated with LLOMe for 2 hours, followed by super-resolution structured illumination microscopy (SR-SIM) imaging of endogenous LAMP1, LGALS3 and M1 poly-Ub. LLOMe induced prominent LMP with accumulation of LAMP1 and M1 poly-Ub at LGALS3-negative and −positive structures in primary mouse cortical neurons **(Figure 6C)**. Image reconstruction in 3D revealed a coat-like distribution of M1 poly-Ub around a subset of LAMP1- and LGALS3-positive lysosomes **(Figure 6D)**, confirming M1 poly-Ub on decorating damaged lysosomes in mouse neurons. Taken together, our data identify M1 poly-Ub as novel cue for lysosomal homeostasis, LMP and degradation of damaged lysosomes, as well as local activation of NF-κB signaling with important implications for inflammation and cell survival.

**Figure 6:**
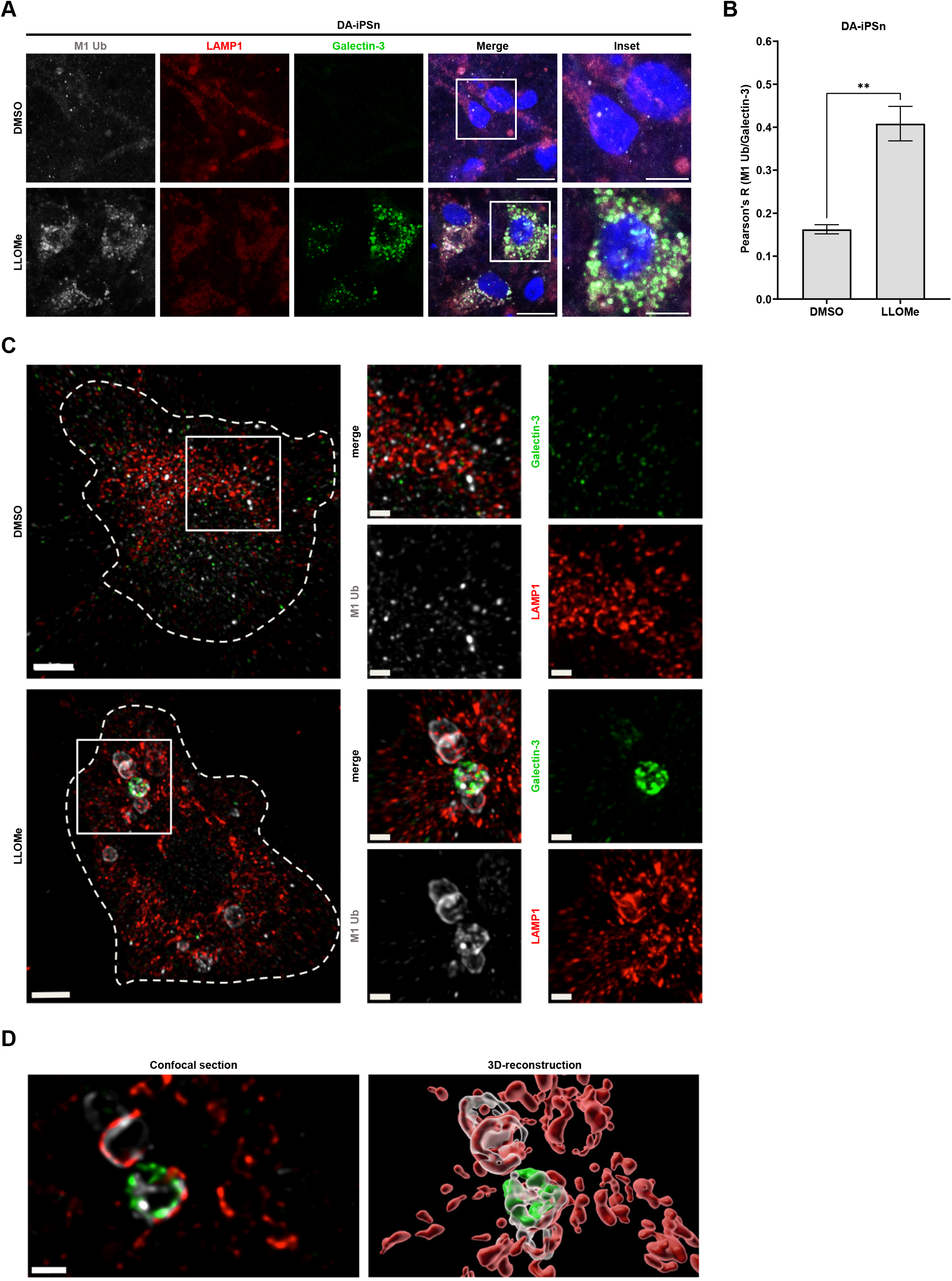
LMP induces M1 poly-Ub of lysosomes in human iPSC-induced dopaminergic neurons and primary mouse cortical neurons. A. Representative immunofluorescence staining of M1 poly-Ub, LAMP1 and LGALS3 in iPSC-derived DA neurons treated with LLOMe (500 µM) for 30 min. Scale bar: 20 µm (10 µm for insets). **B.** Pearson correlation coefficients of M1 poly-Ub/LGALS3 staining from A. Data are shown as mean and SEM from three replicates and at least ten images per replicate. ** p < 0.01. Statistical significance was determined with unpaired *t*-test. **C.** Representative super-resolution structured illumination microscopy (SR-SIM) immunofluorescence staining of untreated and LLOMe treated (2 mM for 2 h) primary cortical mouse neurons (cell soma is outlined) after seven days *in vitro* (DIV) with antibodies against LAMP1 (red), M1 poly-Ub (white) and LGALS3 (green). Scale bars: 3 µm (1 µm for insets). **D.** Precise localization of the antibody signals from C in a single confocal section of a LGALS3 positive lysosome (left) and an Imaris-assisted 3D-reconstruction of the respective area (right). Scale bar: 1 µm.

## Discussion

Lysosomes have diverse pleiotropic functions, ranging from autophagic cargo degradation to membrane repair and cell death. Since LMP is potentially cytotoxic, damaged lysosomes are either repaired or degraded by lysophagy to prevent the release of hydrolytic lysosomal enzymes, such as cathepsins, that mediate cell death. Poly-Ub chains linked via K48 and K63 are deposited on damaged lysosomes and play central roles in lysophagy by recruiting autophagic factors required for degradation, but the role of M1 poly-Ub in lysophagy remains unknown. Here, we reveal that OTULIN negatively regulates the autophagic flux and specifically selects autophagosomal cargo, including the autophagy receptor TAX1BP1 and the glycan-binding LGALS3, implicated in LMP and lysophagy. Loss of OTULIN increases M1 poly-Ub at damaged lysosomes and mediates local NF-κB activation. In addition, our data suggest pro-survival functions of OTULIN-mediated M1 poly-Ub at defect lysosomes. Finally, we show that damaged lysosomes are also modified with M1 poly-Ub in primary mouse neurons and iPSC-derived primary human dopaminergic neurons.

Our findings identify OTULIN as negative regulator of the autophagic flux and specific selection of autophagic cargo. Previous findings suggest that loss of OTULIN promotes autophagy initiation by increasing M1 poly-Ub and stability of ATG13, but has repressive effects on autophagy maturation [34]. This partially correlates with our findings that reveal OTULIN-dependent increases in LC3B lipidation, accumulation of M1 poly-Ub- and LC3B-positive aggregates, degradative compartments in electron microscopy and global and autophagosome-enriched increases in TAX1BP1 and organellar proteins, like LGALS3 and TOMM40, under basal and autophagy-induced conditions. However, we were not able to observe an increased stability of ATG13 upon loss of OTULIN (data not shown), indicating that additional cell- and context-specific mechanisms may control OTULIN-mediated regulation of autophagy.

We reveal that lysosomes are modified with M1 poly-Ub in response to LMP and that OTULIN controls M1 poly-Ub at damaged lysosomes. Up till now, only K48- and K63- linked poly-Ub has been identified at damaged lysosomes that mediate lysophagy [8,11,55]. Ubiquitination of damaged lysosomes occurs in a time-dependent manner. Early upon induction of lysosomal damage, K63-linked poly-Ub serves as important signal for the recruitment of the autophagy machinery [56]. Subsequently, K48 poly-Ub chains peak with some delay compared to K63 poly-Ub at damaged lysosomes and removal of K48 poly-Ub by components of the Endo-Lysosomal Damage response (ELDR), including the AAA-ATPase VCP/p97 and the DUB YOD1, is necessary to drive clearance of damaged lysosomes by lysophagy [8]. Loss of OTULIN only has a minor impact on the clearance of LGALS3-positive damaged lysosomes. However, the observation that M1 poly-Ub co-occurs with K63 poly-Ub might suggest that a combination of both chain types might be required for efficient lysophagy.

We demonstrate that increased M1 poly-Ub at damaged lysosomes locally activates NF-kB signaling via recruitment of NEMO and subsequent phosphorylation-mediated activation of IKK to activate NF-κB-mediated gene transcription. The autophagy- and NF-κB-related functions of M1 poly-Ub at damaged lysosomes correspond with M1 poly-Ub functions in the cell-autonomous recognition of intracellular pathogens [33]. Our observations further revealed that loss of OTULIN induces the phosphorylation-mediated activation of TBK1, correlating with enhanced TBK1 activation described in Otulin^C129A/C129A^Ripk3^−/−^ Casp8^−/−^ bone marrow-derived macrophages (BMDMs) [57]. Interestingly, during TNFα signaling, LUBAC promotes the recruitment and activation of TBK1 and IKKε at TNF receptor 1 signaling complexes (TNFR1-SC) to prevent RIPK1-dependent cell death [58]. TBK1 is also recruited to a mitochondrial NF-κB signaling platform upon TNF receptor activation [59]. Our data suggest similar mechanisms of TBK1 recruitment to LLOMe-damaged lysosomes and local NF-κB activation. This is also supported by previous findings that describe activation of TBK1 at damaged lysosomes and the fact that TBK1 is required for lysophagic flux [31]. Apart from its roles played in lysophagy, TBK1 also regulates other forms of selective autophagy by phosphorylating autophagy receptors such as OPTN, NDP52, TAX1BP1 and p62/SQSTM1 to regulate ubiquitin and LC3 binding [27,29,60,61]. TBK1 activation also plays additional roles in autophagy and affects autophagy progression, both via direct and indirect effects on mTOR inhibition [62–64]. Intriguingly, a potential relation between M1 poly-Ub and mTOR activation has also been revealed in OTULIN-related autoinflammatory syndrome (ORAS), an inflammatory syndrome caused by homozygous mutations in OTULIN. ORAS has been linked to spontaneous and progressive steatotic liver disease and hepatocyte-specific deletion of OTULIN in mice induces steatohepatitis and malignant hepatocellular carcinoma. Intriguingly, liver pathology is associated with increased mTOR activation and rapamycin-mediated mTOR inhibition significantly reduces liver pathology [65].

The deposition of M1 poly-Ub on damaged lysosomes can be prevented by HOIPIN-8-mediated inhibition of HOIP, the catalytic subunit of the M1 poly-Ub-specific E3 ligase complex LUBAC. In addition, lysosomal M1 poly-Ub relies on pre-deposited K63-linked poly-Ub, since inhibition of the dimeric K63-specific E2 enzyme UbcH13/UEV1A E2 completely abrogates M1 poly-Ub accumulation at damaged lysosomes. These findings correspond with an NZF- and K63 poly-Ub-dependent recruitment of HOIP to damaged lysosomes to deposit M1 poly-Ub in a similar manner as LUBAC gets recruited to cytosolic pathogens and death receptor complexes [33,66–68]. Inhibition of HOIP or the K63 poly-Ub-specific UbcH13/UEV1A completely prevented M1 poly-Ub deposition of damaged lysosomes and OTULIN-dependent increases in NF-κB activation. Since K63-linked poly-Ub has been described to occur upstream of NEMO binding to M1 poly-Ub and IKK activation [69,70], we conclude that K63-linked poly-Ub is a prerequisite for M1 poly-Ub. Although the exact identity of M1 poly-Ub substrates during LMP remains unclear, the dependency on K63 poly-Ub supports earlier findings, that describe K63-M1 poly-Ub hybrid chains and K63 poly-Ub as M1 poly-Ub substrates, during activation of immune signaling pathways, such as TNFR or NOD1 [47,48].

The presence of lysosomal proteins in the mass-spectrometry-based autophagosome content profiling indicates that OTULIN might be involved in regulating basal LMP. Inhibition of caspases largely rescued LLOMe-induced cell death, confirming an important role of caspase-dependent apoptosis upon LMP induction, which is in line with reported findings that identify the induction of apoptosis upon lysosomal damage [50,71]. In many cases, LMP-induced cell death is regulated by the release of cathepsins in the cytosol [50]. In addition, cathepsins regulate apoptosis via cleavage of the pro-apoptotic protein Bid [72], the degradation of anti-apoptotic B-cell lymphoma 2 (Bcl-2) family members, like Bcl-2, Bcl-x_L_ and Mcl-1 [73] and the release of cytochrome *c* and caspase activation [74]. Interestingly, our experiments reveal that cathepsin inhibition increased cell death in basal conditions in an OTULIN-dependent manner and even further upon loss of the five major autophagy receptors. These findings indicate that the effects of M1 poly-Ub on lysophagy serve cell-protective functions and prevent cell death. M1 poly-Ub-modified lysosomes might function as cell death platforms upon cathepsin and autophagy inhibition, mediated by M1 poly-Ub in a similar manner as death receptor signaling complexes involved in cell fate checkpoints [67,68].

Our data reveal that induction of LMP in human iPSC-derived dopaminergic neurons triggered accumulation of M1 poly-Ub at damaged lysosomes and formation of LGALS3 puncta at M1 poly-Ub modified lysosomes. This suggests that the response to lysosomal damage is conserved in primary murine and human neurons and reveal the functional relevance of M1 poly-Ub in cellular responses upon lysosomal damage. Neurons are particularly sensitive to lysosomal damage and many neurodegenerative diseases, including Parkinson’s disease, Alzheimer’s disease, amyotrophic lateral sclerosis (ALS) or Niemann-Pick type A, are linked to lysosomal dysfunction and rupture [52,53,75–77]. Thus, a better understanding of the mechanisms that are engaged in the response to lysosomal damage and regulate lysosomal integrity is of utmost importance, especially in neurodegenerative diseases harboring lysosomal dysfunction.

## Supporting information

Scans of original blots

## Acknowledgements

The lab of S.J.L.v.W. received funded by the Deutsche Forschungsgemeinschaft (DFG, German Research Foundation) (WI 5171/1-1, WI 5171/4-1, FU 436/20-1, FU 436/21-1 and project-ID 259130777 – CRC1177), the Deutsche Krebshilfe (70113680), the Wilhelm Sander-Stiftung (2020.008.1), the Frankfurter Stiftung für krebskranke Kinder and the Dr. Eberhard and Hilde Rüdiger Foundation (to S.J.L.v.W.). The work from F.Z. is supported by the DFG (ZU 409/1-1 – RTG 2162) and the Interdisciplinary Center for Clinical Research (IZKF) at the University Hospital of the University of Erlangen-Nuremberg (Jochen-Kalden funding programme N8). C.B. was supported by the DFG within the frameworks of the Munich Cluster for Systems Neurology (EXC 2145 SyNergy – ID 390857198 (C.B.), the Collaborative Research Center (CRC) 1177 (ID 259130777) and the project grant BE4685/8-1. M.M is supported by the ALW Open Programme (ALWOP.355). K.F.W. is funded by the DFG (WI/2111-6, WI/2111-8, FOR 2848) and Germany’s Excellence Strategy - EXC 2033 - 390677874 – RESOLV and by the Michael J. Fox Foundation (Grant ID: 021968). SR-SIM microscopy was funded by the DFG and the State Government of North Rhine-Westphalia (INST 213/840-1 FUGG). The authors acknowledge C. Hugenberg for critical reading of the manuscript.

## Author contributions

L.Z. and M.D. performed most of the experiments. S.Z. and C.B. performed and analyzed the APEX2-based autophagosome content profiling. D.B., J.V. and F.Z. cultured iPSCs, differentiated DA-iPSn and performed immunofluorescence staining and imaging of DA-iPSn. V.B. and K.F.W. performed experiments and data analysis with primary mouse neurons and SR-SIM. M.C.M. conducted electron microscopy and quantification of the EM images. N.W. generated cell lines. L.Z., M.D. and S.J.L.v.W. analyzed the data, prepared the manuscript and designed the study. All authors read, commented and discussed the manuscript.

## Materials and methods

### Cell lines and chemicals

The human GBM cell line MZ-54 [78,79], human embryonic kidney cells (HEK293T®, ATCC CRL-3216™) and HeLa cells (WT and Penta KO cells were gifts from Richard Youle [43]) were cultured in DMEM + GlutaMAX (Thermo Fisher Scientific) supplemented with 1% penicillin-streptomycin (Thermo Fisher Scientific) and 10% fetal bovine serum (FBS, Thermo Fisher Scientific) at 37 °C and 5% carbon dioxide (CO_2_) in a 95% humidified atmosphere. MZ-54 and HEK293T cells were regularly tested for mycoplasma infection and authenticated by the Leibniz Institute DSMZ GmbH (Braunschweig, Germany). Cell line authentication was performed using DNA profiling with different highly polymorphic short tandem repeats (STR) loci.

### iPSC culture and neuronal differentiation

Isogenic control Parkinson disease patient-derived human (patient #1) induced pluripotent stem cells (iPSC), carrying a triplication in the alpha-synuclein gene, were corrected with CRISPR/Cas9 gene editing (isogenic control of 3×1) and were previously characterized extensively [80,81]. iPSCs were cultivated in mTeSR1 plus media (Stemcell, 100-0276) on Matrigel (Corning, 354234)-coated 6-well plates and differentiated into dopaminergic (DA) neurons using an established protocol [82] at a constant of 37 °C and 5% CO_2_. Between day 25-30 of differentiation, neurons were seeded on poly-D-lysine (Merck, P1149) and LAM/Laminin (Merck, 11243217001) coated 24-well plates at a number of 3×10^5^ per well on 12-mm cover glasses for immunofluorescence (or at 4×10^5^ for Western blot). Neurons were maintained in Neurobasal media (Thermo Fisher Scientific, 21103049) containing NeuroCult SM1 Neuronal Supplement (Stemcell, 05711) and 1 % penicillin-streptomycin until ∼day 90, when neurons were treated for 30 min with LLOMe (500 µM) or the equal volume of DMSO as control.

### Primary mouse cortical neuronal cultures

Mouse cortices from embryonic day 18 (E18) embryos were dissected into ice-cold HBSS and dissociated in acetylated trypsin (Sigma) for 20 min at 37°C. Subsequently, tissues were rinsed with HBSS and treated with DNase I to break down DNA and triturated with a cell strainer. The cell suspension was resuspended in Neurobasal medium (Invitrogen) and plated in 24-well plates on coverslips (Laboratory Glassware Marienfeld) coated with poly-L-lysine (Sigma) and laminin (Sigma) at a concentration of 50,000 neurons/well for immunocytochemistry. Medium was replaced after 45 min. Primary neurons were cultured in Neurobasal medium with B27 (Invitrogen) and glutamine (Invitrogen) in a humidified incubator (37°C, 5% CO_2_). At day 7 *in vitro*, cortical neurons were treated with 2 mM LLOMe for 2 h and fixed in 4% paraformaldehyde in PBS for 15 min and washed with PBS. Protocols were performed in compliance with institutional and governmental regulations.

### Drug treatments

Bafilomycin A1 (BafA1) (Selleckchem, S1413), loperamide (LOP) hydrochloride (Enzo Life Sciences, ALX-550-253-G005), torin 1 (Selleckchem, S2827), cycloheximide (CHX) (Merck, C7698), L-leucyl-L-leucine methyl ester (LLOMe) (Cayman Chemical, 16008), HOIPIN-8 (MedChemExpress, HY-122882), zVAD.fmk (Bachem, 4027403), TPCA-1 (Selleckchem, S2824), NSC697923 (Selleckchem, S7142) and Pepstatin A (Sigma-Aldrich, EI10) were used as described in the figure legends.

### Generation of OTULIN KO cell lines

Generation of CRISPR/Cas9-mediated MZ-54 and HeLa OTULIN KO was done as previously described [83] using guide RNAs against human OTULIN (5’-AAGAGTCCCTTACTCTGCTG-3’, 5’-ACCACGGACTCGCCGTATGG-3’, 5’-ATTGCTTATACATGAAAGAG-3’, 5’-TGAACTATTCACAAATGAGG-3’), GFP as non-human target (NHT) control (5’-GGAGCGCACCATCTTCTTCA-3’, 5’-GCCACAAGTTCAGCGTGTC-3’, and 5’-GGGCGAGGAGCTGTTCACCG-3’) cloned into pLentiCRISPRv2 (Addgene, 52961, deposited by Feng Zhang). To generate lentiviral particles, HEK293T cells were co-transfected with pLenti-CRISPRv2 plasmids and the packaging plasmids pPAX2 (Addgene #12260, deposited by Didier Trono) and pMD2.G (Addgene #12559, deposited by Didier Trono) using FuGENE® HD Transfection Reagent (Promega, E2311) at a ratio of 1:3 (µg DNA/µl FuGENE) according to the manufacturer’s instructions in antibiotic-free medium. 24 h after transfection the medium was changed and the viral supernatant was harvested 48 and 72 h after transfection, pooled and sterile-filtered through a 0.45 µm filter (MerckMillipore, Darmstadt, Germany). Target cells were transduced with 0.5 mL of viral supernatant in 3 mL of total medium supplemented with 8 µg/mL polybrene (Merck Sigma). 2 to 4 days post transduction, cells were reseeded in selection medium containing 1 µg/mL puromycin. Successful depletion of OTULIN in the bulk culture was confirmed by Western blot analysis.

### Generation of myc-APEX2-LC3B-expressing MZ-54 cells

To stably generate myc-APEX2-LC3B-expressing cells, HEK293T cells were transfected with pHAGE-myc-APEX2-LC3B [39] and the packaging plasmids pHDM-VSV-G, pHDM-Tat1b, pHDM-Hgpm2 and pRC-CMV-Rev1b using Lipofectamine 2000. pHDM-VSV-G (Addgene #164440; RRID:Addgene_164440), pHDM-Tat1b (Addgene plasmid #164442; RRID:Addgene_164442), pHDM-Hgpm2 (Addgene plasmid #164441; RRID:Addgene_164441) and pRC-CMV-Rev1b (Addgene plasmid #164443, RRID:Addgene_164443) were gifts from Alejandro Balazs.

MZ-54 cells were transduced with lentiviral particles containing pHAGE-myc-APEX2-LC3B plasmids in medium with 8 µg/mL polybrene. 2 to 4 days post transduction, selection was started with 2 µg/mL puromycin (pHAGE-myc-APEX2-LC3B). Successful protein expression was verified by Western blot analysis.

### RNAi-mediated silencing of OTULIN

RNAi-mediated silencing of OTULIN expression in HeLa cells was done with the following three Silencer™ Select siRNAs (Thermo Fisher Scientific): siOTULIN#1, s40306; siOTULIN#2, s40307; siOTULIN#3, s40308 (20 nM each, 60 nM in total) and a Silencer™ Select negative control siRNA (Thermo Fisher Scientific, #4390843, 60 nM). For reverse transfection, siRNA-lipid complexes were prepared in OptiMEM (Thermo Fisher Scientific) with Lipofectamine™ RNAiMAX Transfection Reagent (Thermo Fisher Scientific) according to the manufacturer’s instructions and incubation for 5 min at room temperature (RT). The reaction mixture was distributed dropwise in cell culture plates and cells in antibiotic-free medium were seeded on top. 24 h after transfection, medium was changed and cells were reseeded for experiments 48 h after transfection. Knockdown efficiency was confirmed by Western blot analysis.

### Determination of cell death

Cell death was measured by PI (Sigma-Aldrich, P4864) and Hoechst33342 (Sigma-Aldrich, 14533) (working concentrations: 10 µg/mL Hoe and 1 µg PI) double staining and fluorescence-based analysis of dead (PI-positive) cells in relation to the total cell number (Hoechst-positive cells) with the ImageXpress® Micro XLS Widefield High-Content Analysis System and MetaXpress® Software (Molecular Devices Sunnyvale, CA, USA).

### Immunofluorescence analysis

For immunofluorescence analysis, cells were seeded on microscope coverslips into Greiner 12-well plates with a density of 30,000 cells per well. After treatment, cells were washed with PBS and fixed with 4% formaldehyde (FA, 28908) for 5 min at RT. Cells were washed again with PBS, followed by blocking and permeabilization in permeabilization buffer (PB) (0.1% Saponin (Carl Roth, 9622.1) and 5 mM MgCl_2_ (Carl Roth, 2189.1) in PBS) supplemented with 10% FBS for 1 h at RT. First antibody incubation was performed overnight (ON) at 4 °C in PB with 5% FBS. The following primary antibodies were used for immunofluorescence staining: mouse anti-LAMP1 (DSHB Cat# H4A3, RRID:AB_2296838, 1:200), rabbit anti-LAMP1 (D2D11) XP (Cell Signaling Technology Cat# 9091, RRID:AB_2687579, 1:200), mouse anti-LC3 Clone 4E12 (MBL International Cat# M152-3, RRID:AB_1279144, 1:100), rat anti-mouse/human Galectin 3 (BioLegend Cat# 125402, RRID:AB_1134238, 1:100), rabbit anti-TAX1BP1 (Sigma-Aldrich Cat# HPA024432, RRID:AB_1857783, 1:100), rabbit anti-NAK (phospho S172) (Abcam Cat# ab109272, RRID:AB_10862438, 1:100), rabbit anti-IKBKG (Sigma-Aldrich Cat# HPA000426, RRID:AB_1851572, 1:100), rabbit anti-phospho-IKKα/β (Ser176/180) (16A6) (Cell Signaling Technology Cat# 2697, RRID:AB_2079382, 1:100), rabbit anti-RNF31/HOIP (Abcam Cat# ab46322, RRID:AB_945269, 1:100), rabbit anti-Ub K63-specific, clone Apu3 (Millipore Cat# 05-1308, RRID:AB_1587580, 1:100) and human anti-M1 Ub (Genentech, 1:2000). After 3 washing steps with PB for 5 min at RT, secondary antibodies diluted 1:500 in PB with 5% FBS were added for 2 h at RT. Secondary antibodies for immunofluorescence were goat anti-mouse IgG (H+L) Alexa Fluor™ 555 (Thermo Fisher Scientific Cat# A-21422, RRID:AB_2535844), goat anti-rabbit IgG (H+L) Alexa Fluor™ 555 (Thermo Fisher Scientific Cat# A-21428, RRID:AB_2535849), goat anti-mouse IgG (H+L) Alexa Fluor™ 647 (Thermo Fisher Scientific Cat# A-21235, RRID:AB_2535804), goat anti-rabbit IgG (H+L) Alexa Fluor™ 647 (Thermo Fisher Scientific Cat# A-21244, RRID:AB_2535812), goat anti-human IgG (H+L) Alexa Fluor™ 647 (Thermo Fisher Scientific Cat# A-21445, RRID:AB_2535862) and donkey anti-rat IgG (H+L) Alexa Fluor™ 488 (Thermo Fisher Scientific Cat# A-21208, RRID:AB_2535794). After 2 washing steps with PB coverslips were rinsed in ddH_2_O and mounted on glass slides with ProLong™ Diamond Antifade Mountant with DAPI (Thermo Fisher Scientific, P36966). Image acquisition was performed with the Leica SP8 laser-scanning microscope (LSM) (Leica Microsystems, Wetzlar, Germany) using 63× magnification and Type F Immersion Oil (Thorlabs, MOIL-10LF). For immunofluorescence of DA-iPSn, cells were washed with PBS after removal of the secondary antibodies and stained with DAPI (Sigma-Aldrich, Cat#D8417) diluted 1:10,000 in PBS for 10 min at RT. After two washing steps with PBS and one washing step with ddH_2_O, DA-iPSn were mounted on glass slides with ProLong™ Gold Antifade Mountant (Thermo Fisher Scientific, Cat# P36930). Image acquisition of DA-iPSn was performed with a Zeiss LSM 780 (Carl Zeiss Microscopy GmbH) and the Zeiss blue software (ZEN lite 2012). For immunofluorescence on primary mouse neurons, neurons were permeabilized and blocked in 0.1% (v/v) Triton X-100, 5% (v/v) goat serum, 1% BSA (w/v) in PBS for 1 h. Cells were stained with primary antibodies against LGALS3 (1:250) (Thermo Fisher Scientific Cat# 14-5301-82, RRID:AB_837132), LAMP1 (1:250) (Abcam Cat# ab24170, RRID:AB_775978) and M1-poly Ub 1:1000 (1E3, Genentech) in 0.1% (v/v) Triton X-100, 5% (v/v) goat serum, 1% BSA (w/v) in PBS at 4°C overnight, washed 3 times with PBS, and incubated with fluorescent dye-conjugated secondary antibodies Alexa Fluor 488, 555, or 647 (Thermo Scientific), at a dilution of 1:1000 for 1 h at RT. After 3 washing steps in PBS, cells were incubated with DAPI (Sigma) and mounted in EverBrite Hardset Mounting medium (Biotium). Super-resolution structured illumination microscopy (SR-SIM) was performed using a Zeiss Elyra7 equipped with an 63× NA 1.4 OIL objective and the ZEN image acquisition software including the SIM2 upgrade (Zeiss, Oberkochen, Germany). 3D image reconstruction was performed using the surface function of the Imaris 10.0.1 image analysis software (Bitplane, Zürich, Switzerland). Additional image analysis was performed with ImageJ (version 1.53u) and FIJI (Fiji Is Just ImageJ; version 2.14.0/1.54f). Quantification of puncta was done manually or using the FIJI plugin Aggrecount (v1.13) with at least three independent microscopic views. Pearson’s correlation coefficient was calculated with the FIJI plugin Coloc2.

### Western blot analysis

Cell lysis was performed using RIPA lysis buffer (50 mM Tris-HCl [Carl Roth, 9090.3], pH 8, 150 mM sodium chloride, 1% IGEPAL CA-630/NP-40 [Sigma-Aldrich, 56741], 2 mM magnesium chloride, 0.5% sodium deoxycholate) supplemented with protease inhibitor complex (PIC) (Roche, Grenzach, Germany), 0.1% SDS, 1 mM sodium orthovanadate, 5 mM sodium fluoride, 1 mM phenylmethylsulfonyl fluoride and Pierce Universal Nuclease (Thermo Fisher Scientific, 88701). Cells were lysed with RIPA lysis buffer for 20 min on ice, followed by centrifugation at 18,000xg and 4 °C for 25 min and measurement of the protein concentration using the Pierce™ BCA Protein Assay Kit according to the manufacturer’s protocol. Cell lysates were boiled in 6x SDS loading buffer (350 mM Tris Base pH 6.8, 38% glycerol, 10% SDS, 93 mg/ml dithiothreitol (DTT), 120 mg/ml bromophenol blue), loaded on self-cast polyacrylamide gels (gel percentages 10 or 13.5%) and subjected to SDS-polyacrylamide gel electrophoresis (SDS-PAGE). SDS-PAGE was followed by transfer of proteins on a nitrocellulose membrane with a semi-dry blotting system (BioRad) and protein expression was analyzed by incubation with the following primary antibodies: human anti-M1 Ub (Genentech, 1:2000), rabbit anti-OTULIN (Cell Signaling Technology Cat# 14127, RRID:AB_2576213,1:1000), rabbit anti-RNF31/HOIP (Abcam Cat# ab46322, RRID:AB_945269, 1:1000), mouse anti-HOIL-1 clone 2E2 (Millipore Cat# MABC576, RRID:AB_2737058, 1:1000), rabbit anti-Sharpin (D4P5B) (Cell Signaling Technology Cat# 12541, RRID:AB_2797949, 1:1000), rabbit anti-LC3B (Thermo Fisher Scientific Cat# PA1-16930, RRID:AB_2281384, 1:2000), mouse anti-Vinculin clone hVin-1 (Sigma-Aldrich Cat# V9131, RRID:AB_477629, 1:2000), mouse anti-GAPDH (Hytest Cat# 5G4cc-6C5cc, RRID:AB_2858176, 1:5000), mouse anti-β-Actin (Sigma-Aldrich Cat# A5441, RRID:AB_476744, 1:10,000), rabbit anti-TAX1BP1 (D1D5) (Cell

Signaling Technology Cat# 5105, RRID:AB_11178939, 1:1000), rabbit anti-p62 (MBL International Cat# PM045, RRID:AB_1279301, 1:5000), mouse anti-Galectin 3 (Thermo Fisher Scientific Cat# MA1-940, RRID:AB_2136775, 1:1000), rabbit anti-TOMM40 [EPR6932(2)] (Abcam, ab185543, 1:1000), rabbit anti-NAK (phospho S172) (Abcam Cat# ab109272, RRID:AB_10862438, 1:1000), rabbit anti-TBK1 (Cell Signaling Technology Cat# 38066, RRID:AB_2827657, 1:1000), rabbit anti-Cathepsin D (E179) (Cell Signaling Technology Cat# 69854, 1:1000), goat anti-Biotin (Thermo Fisher Scientific Cat# 31852, RRID:AB_228243, 1:10000), rabbit anti-KBTBD7 (Novus Cat# NBP1-92040, RRID:AB_11024468, 1:1000), goat anti-Myc Tag (Bethyl Laboratories Cat# A190-104A, RRID:AB_66864, 1:1000), rabbit anti-LAMP1 (D2D11) XP (Cell Signaling Technology Cat# 9091, RRID:AB_2687579, 1:1000), mouse anti-Neurofilament (Biolegend, Cat# 837904, RRID:AB_2566782, 1:500), mouse anti-Tyrosin hydroxylase (Sigma-Aldrich, Cat# T1299, RRID:AB_477560, 1:500), rabbit anti-Synapsin I (Innovative Research, Cat# A6442, RRID:AB_1502084, 1:500) and mouse anti-tubulin beta-3 (BioLegend, Cat# 801202, RRID:AB_10063408, 1:500). All primary antibodies were incubated overnight at 4°C, except for the human anti-M1 Ub antibody which was incubated for 1 h at RT. Secondary antibody incubation was done for 1 h at RT with the following horseradish peroxidase (HRP)-coupled antibodies: HRP-conjugated goat anti-mouse IgG (Abcam Cat# ab6789, RRID:AB_955439), HRP-conjugated goat anti-rabbit IgG (Abcam Cat# ab6721, RRID:AB_955447) and HRP-conjugated goat anti-human IgG (AP309P, Merck KGaA). Protein detection was done with enhanced chemiluminescence (ECL) using Pierce™ ECL Western Blotting-Substrate (Thermo Fisher Scientific). For stripping of membranes, 0.4 M NaOH was used. Representative blots of at least two independent experiments are shown. If the samples of one experiment are detected on multiple Western blotting membranes, only one representative loading control is shown for clarity.

### Proximity labeling, proteinase K digest and streptavidin pulldown

APEX2-LC3B proximity labeling, proteinase K digest and streptavidin pulldown followed by mass spectrometry was performed as described before [39]. In brief, APEX2-mediated biotinylation of cells was done by incubation with 500 µM biotin-phenol for 30 min at 37 °C, followed by addition of 1 mM H_2_0_2_ for 1 min at RT. The reaction was stopped with quencher solution (1 mM sodium azide, 10 mM sodium ascorbate and 5 mM Trolox in DPBS) before washing with DPBS and harvesting of cells. For proteinase K digest, cells were resuspended in homogenization buffer I (10 mM KCl, 1.5 mM MgCl_2_, 10mM HEPES-KOH and 1 mM DTT pH 7.5), incubated on an overhead shaker and dounce homogenized. Homogenization buffer (375 mM KCl, 22.5 mM MgCl_2_, 220 mM HEPES-KOH and 0.5 mM DTT, pH 7.5) was added and lysates were cleared by centrifugation at 600xg for 10 min. For immunoblotting, 30 µg/mL proteinase K was added to the samples for 30 min with or without 0.2% Triton-X-100 as control. Digestion was then stopped with PMSF, samples were supplemented with 6x SDS loading buffer (composition see above) and subjected to SDS-PAGE and immunoblotting. For mass spectrometry, samples were incubated with 100 µg/mL proteinase K for 1 h and the control samples were additionally supplemented with 0.1% RAPIGest™. To separate digested from membrane-protected material, centrifugation at 17,000xg for 15 min was performed. For streptavidin pulldown, biotin-labeled pellets were resuspended in RIPA buffer with quenching components (50 mM Tris, 150 mM NaCl, 0.1% SDS, 0.5% sodium deoxycholate, 1% Triton X-100, 1x protease inhibitors (Roche), 1x PhosStop (Roche), 1 mM sodium azide, 10 mM sodium ascorbate and 1 mM Trolox), sonified and centrifuged at 10,000xg. The supernatants were applied on pre-equilibrated Streptavidin-Agarose resins and incubated overnight. Next, samples were washed 3 times in RIPA buffer with quenching components and 3 times in 3 M Urea buffer (in 50 mM NH_4_HCO_3_) before incubation with 5 mM TCEP for 30 min at 55 °C and shaking. To alkylate samples, 10 mM IAA was added for 20 min at RT and quenched by addition of 20 mM DTT. After 2 washing steps with 2 M Urea buffer (in 50 mM NH_4_HCO_3_) over-night trypsin digestion was performed with 1 µg trypsin per 20 µl beads at 37 °C. Supernatants were collected from the resin, washed twice with 2 M Urea buffer and acidified with 1% trifluoroacetic acid. The sample volume was then decreased by vacuum centrifugation and digested peptides were desalted on custom-made C18 stage tips. Finally, samples were reconstituted with 0.5% acetic acid for mass spectrometric analysis.

### Mass spectrometry data collection and analysis

Samples were loaded onto 75 µm x 15 cm fused silica capillaries (custom-made) packed with C18AQ resin (Reprosil-Pur 120, 1.9 µm, Dr. Maisch HPLC) using an Easy-nLC1200 liquid chromatography. Peptide mixtures were separated at a 400 nl/min flow rate using a 35 min ACN gradient in 0.5% acetic acid (5–38% ACN gradient for 23 min followed by 38-60% ACN gradient for 3 min and 60-95% ACN gradient for 2 min plus another 3 min at 95% CAN prior to 95%-5% gradient for 2 min and another 2 min at 5% ACN) and detected on an Q Exactive HF mass spectrometer (Thermo Scientific). Dynamic exclusion was enabled for 20 sec and singly charged species, charge states above 8 or species for which a charge could not be assigned were rejected. Mass spectrometry raw data were analyzed using MaxQuant (version 1.6.0.1) [84] and a human Uniprot FASTA reference proteome (UP000005640) in reverted decoy mode with the following allowance: methionine oxidation and protein N-terminus acetylation as variable modifications, cysteine carbamidomethylation as fixed modifications, 2 missed cleavages and 5 modifications per peptide, minimum peptide length of 7 amino acids, first search peptide tolerance of ±20 ppm, main search peptide tolerance of ±4.5, match between runs, label-free quantification (LFQ), as well as protein, peptide and site level false discovery rates of 0.01. For further processing, MaxQuant output files (protein groups) were loaded into Perseus (version 1.6.5.0) [85] where matches to common contaminants, reverse identifications, identifications based only on site-specific modifications and with less than 2 peptides and MS/MS counts were removed. Identifications from the RAPIGest™ and proteinase K treated controls were subtracted from proteinase K treated APEX2-LC3B samples. Only proteins with LFQ intensities in 2 out of 3 biological replicates in at least one experimental group were kept for the subsequent label-free quantification. LFQ intensities were log2 transformed and missing values were replaced with random numbers drawn from a normal distribution. Student *t*-tests were used to determine the statistical significance of the abundance alterations of proteins identified in the proximity of APEX2-LC3B between experimental conditions.

### Electron microscopy

For conventional transmission electron microscopy, MZ-54 NHT and OTULIN KO GBM cells were treated with LOP (17.5 µM). After 16 h, cells were prepared for electron microscopy as previously described [86]. Cell sections were analyzed using a Talos™ F200i TEM (Thermo Fisher Scientific, Eindhoven, The Netherlands). The average number of autophagosomes and degradative compartments (amphisomes, lysosomes and autolysosomes) per cell section was determined by counting these compartments through 120 cell sections per condition, randomly selected from three independent grids.

### RNA isolation, cDNA synthesis and quantitative real-time PCR (qRT-PCR)

For analysis of gene expression, cells were seeded in 6-well plates at 100,000 cells per well. Total RNA from cells was isolated using the peqGOLD total RNA isolation kit (VWR, Radnor, PA, USA) according to the manufacturer’s protocol and RNA concentrations of the samples were determined with a NanoDrop 1,000 spectrometer (VWR, Darmstadt, Germany). 1 µg of RNA per sample was used for cDNA synthesis with the RevertAid H Minus First Strand Kit (ThermoFisher Scientific) according to the manufacturer’s protocol. To determine gene expression levels, SYBR green-based qRT-PCR was performed using the SYBR™ Green PCR Master Mix (Thermo Fisher Scientific, #4312704) and the 7900GR Fast Real-time PCR system (Applied Biosystems, Darmstadt, Germany). Relative gene expression levels were calculated using the 2−ΔΔCT-method [87] and normalized to the housekeeping gene *RPII*. For quantification and statistical analysis, all experiments were performed at least three times and in technical triplicates. All primers used in this study were purchased by Eurofins (Hamburg, Germany) and were as follows: *IL6*: forward: 5’-GATGAGTACAAAAGTCCTGATCCA-3’, reverse: 5’-CTGCAGCCACTGGTTCTGT-3’, *IL8*: forward: 5’-CTCTTGGCAGCCTTCCTGATT-3’, reverse: 5’-TATGCACTGACATCTAAGTTCTTTAGCA-3’, *TNFA*: forward: 5’-ACAACCCTCAGACGCCACAT-3’, reverse: 5’-TCCTTTCCAGGGGAGAGAGG-3’, *RPII*: forward: 5’-GCACCACGTCCAATGACAT-3’, reverse: 5’-GTGCGGCTGCTTCCATAA-3’.

### Statistical analysis

Results are presented as mean +/- SEM. Statistical analysis was performed with GraphPad Prism 9 Software (GraphPad Software, La Jolla CA, USA) using *t*-test or two-way analysis of variance (ANOVA) followed by Tukey’s multiple comparisons tests. *p*-values were interpreted as follows: * p < 0.05, ** p < 0.01, *** p < 0.001, **** p <0.0001, ns: not significant.

## Supplemental information

**Supplemental Figure 1 related to Figure 1. A.** Western blot analysis of MYC and OTULIN expression in myc-APEX2-LC3B-expressing MZ-54 NHT and OTULIN KO GBM cells to confirm myc-APEX2-LC3B overexpression. Vinculin was used as loading control. Representative blots of at least two independent experiments are shown. **B.** Western blot analysis of Biotin, p62, MYC, OTULIN and KBTBD7 expression in myc-APEX2-LC3B-expressing MZ-54 NHT and OTULIN KO GBM cells, treated with BafA1 (200 nM) for 2 h, homogenized and incubated with Proteinase K, Triton X-100 or both. KBTBD7 was used as negative control. Representative blots of at least two independent experiments are shown. **C.** Western blot analysis of Biotin, MYC, OTULIN and LC3B expression in myc-APEX2-LC3B-expressing MZ-54 NHT and OTULIN KO GBM cells, treated with LOP (17.5 µM) for 16 h (left) or BafA1 (200 nM) for 2 h (right). GAPDH was used as loading control. Representative blots of at least two independent experiments are shown. **D.** Western blot analysis of TAX1BP1, p62, OTULIN, TOMM40 and LGALS3 expression in MZ-54 NHT and OTULIN KO GBM cells, treated with BafA1 (200 nM) for 2 h, homogenized and incubated with Proteinase K, Triton X-100 or both. KBTBD7 was used as negative control. Representative blots of at least two independent experiments are shown. **E.** Western blot analysis of TAX1BP1 and OTULIN expression in MZ-54 NHT and OTULIN KO GBM cells, treated with cycloheximide (CHX, 10 µg/ml) for the indicated time points. GAPDH was used as loading control. Representative blots of at least two independent experiments are shown. **F.** Idem as E, but MZ-54 NHT and OTULIN KO GBM cells were treated with CHX (10 µg/ml) and BafA1 (200 nM) for 8 and 24 h. GAPDH was used as loading control. Representative blots of at least two independent experiments are shown.

**Supplemental Figure 2 related to Figure 2. A.** Representative immunofluorescence staining of M1 poly-Ub/TAX1BP1 (upper panel) and LAMP1/TAX1BP1 (lower panel) in MZ-54 NHT and OTULIN KO GBM cells treated with LLOMe (500 µM) for 16 h. Scale bar: 20 µm (10 µm for insets). **B.** Idem as A, but after treatment of MZ-54 NHT and OTULIN KO GBM cells with LLOMe (500 µM) for 2 h, LLOMe was washed out and cells were incubated in fresh medium for additional 4 h and stained for LAMP1 and LGALS3. Scale bar: 20 µm (10 µm for insets). **C.** Quantification of LGALS3 puncta. Mean and SEM of 38 cells from three independent experiments are shown. * p < 0.05, ** p < 0.01, ns: not significant. **D.** Western blot analysis of p-TBK1 (S172), TBK1 and OTULIN expression in MZ-54 NHT and OTULIN KO GBM cells treated with LLOMe (500 µM) for 2 h. GAPDH was used as loading control. Representative blots of at least two independent experiments are shown. **E.** Western blot analysis of p-TBK1 (S172), TBK1, OTULIN and M1 poly-Ub expression in MZ-54 NHT and OTULIN KO GBM cells pre-treated with HOIPIN-8 (30 µM) for 24 h and treated with LLOMe (500 µM) for 2 h. β-Actin was used as loading control. Representative blots of at least two independent experiments are shown. **F.** Representative immunofluorescence staining of LAMP1/Galectin-3 in HeLa WT and OTULIN KO cells treated with LLOMe (500 µM) for 2 h. Scale bar: 20 µm (10 µm for insets). **G.** Idem as F, but HeLa WT and OTULIN KO cells treated with LLOMe (500 µM) for 2 h were stained for M1 poly-Ub/LAMP1. Scale bar: 20 µm (10 µm for insets). Statistical significance in (**C**) was determined with two-way analysis of variance (ANOVA) followed by Tukey’s multiple comparisons tests.

**Supplemental Figure 3 related to Figure 3. A.** Quantification of M1 poly-Ub- and pIKKα/β (Ser176/180)-positive puncta (left panel) and pIKKα/β (Ser176/180)- and LAMP1-positive puncta (right panel) of MZ-54 NHT and OTULIN KO GBM cells treated with LLOMe (500 µM) for 2 h. Mean and SEM of 21-25 cells from at least four independent microscopic views are shown. **** p < 0.0001. **B.** Representative immunofluorescence staining of M1 poly-Ub/NEMO and M1 poly-Ub/ pIKKα/β (Ser176/180) in HeLa WT and OTULIN KO cells treated with LLOMe (500 µM) for 2 h. Scale bar: 20 µm (10 µm for insets). **C.** Quantification of M1 poly-Ub- and NEMO-positive puncta in MZ-54 OTULIN KO GBM cells pre-treated with HOIPIN-8 (30 µM) for 1 h before addition of LLOMe (500 µM) for 2 h. Mean and SEM of 36 cells from at least five independent microscopic views are shown. **** p < 0.0001. **D.** Quantification of M1 poly-Ub- and pIKKα/β (Ser176/180)-positive puncta of MZ-54 NHT and OTULIN KO GBM cells pre-treated with TPCA-1 (5 µM) for 1 h before addition of LLOMe (500 µM) for 2 h. Mean and SEM of 31 cells from at least six independent microscopic views are shown. **** p < 0.0001. Statistical significance was determined with two-way analysis of variance (ANOVA) followed by Tukey’s multiple comparisons tests (**A**) and with student’s *t*-test (**C&D**).

**Supplemental Figure 4 related to Figure 6.** Western blot analysis of neurofilament, synapsin-I, tyrosine hydroxylase (TH) and β3-Tubulin expression in iPSC-derived DA neurons treated with LLOMe (500 µM) for 30 min. The isogenic, CRISPR-corrected control of patient-derived human induced pluripotent stem cells (iPSCs) (patient #1), carrying a triplication in the alpha-synuclein gene, is shown (Isogenic control of 3×1). GAPDH was used as loading control.

