## Supplementary figures and images for "Linear ubiquitination at damaged lysosomes induces local NF-κB activation and controls cell survival"

### Scans of original blots

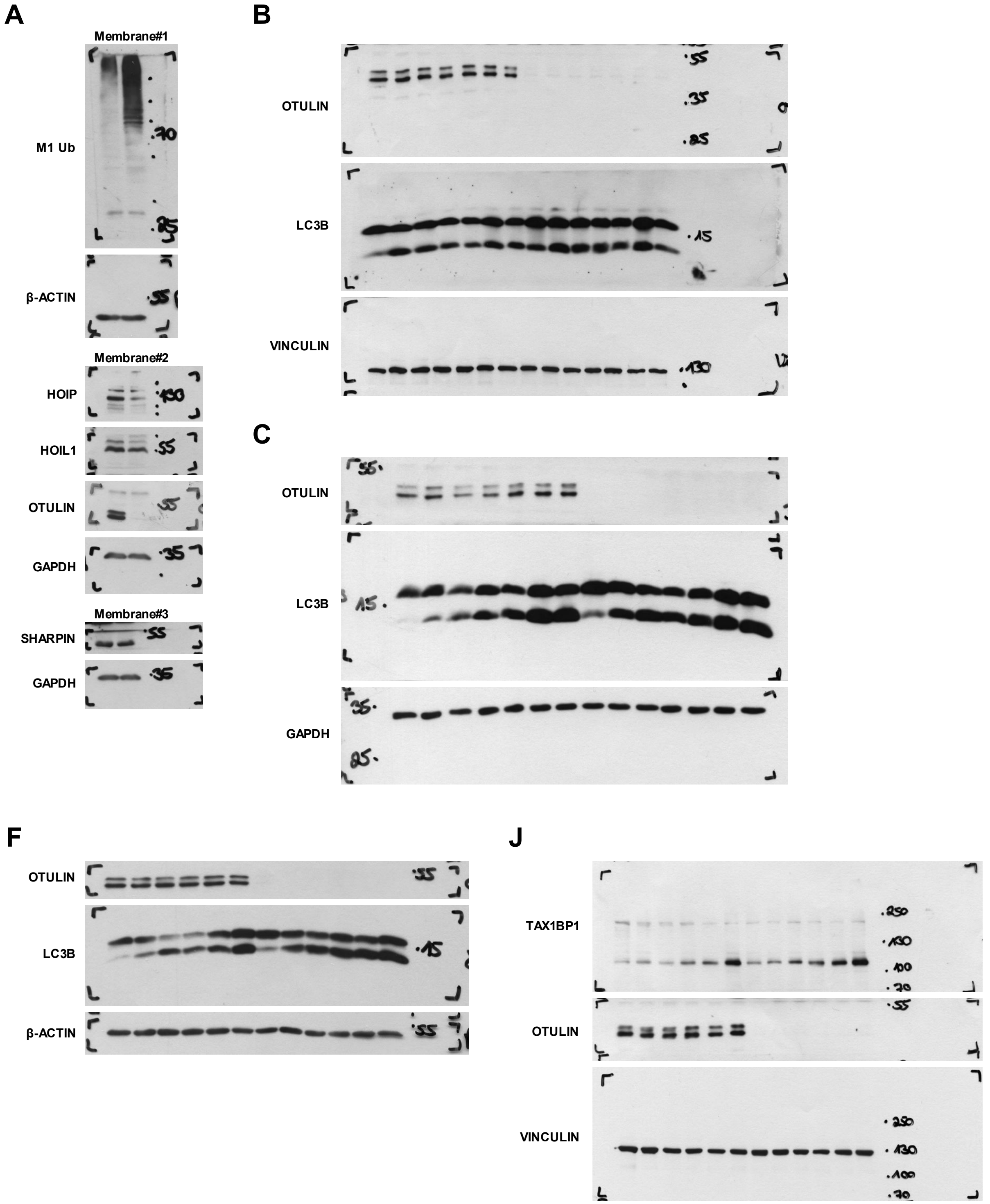

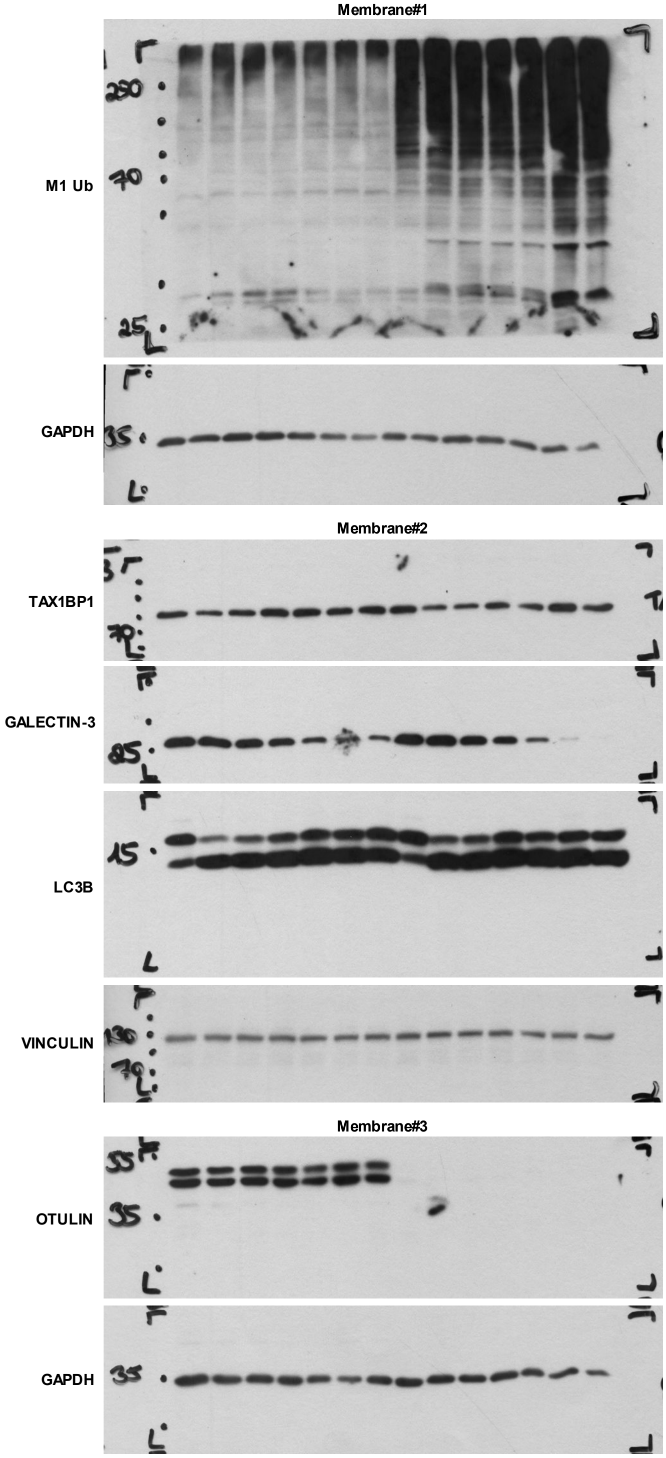

C

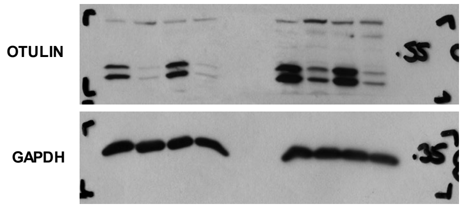


Figure 5

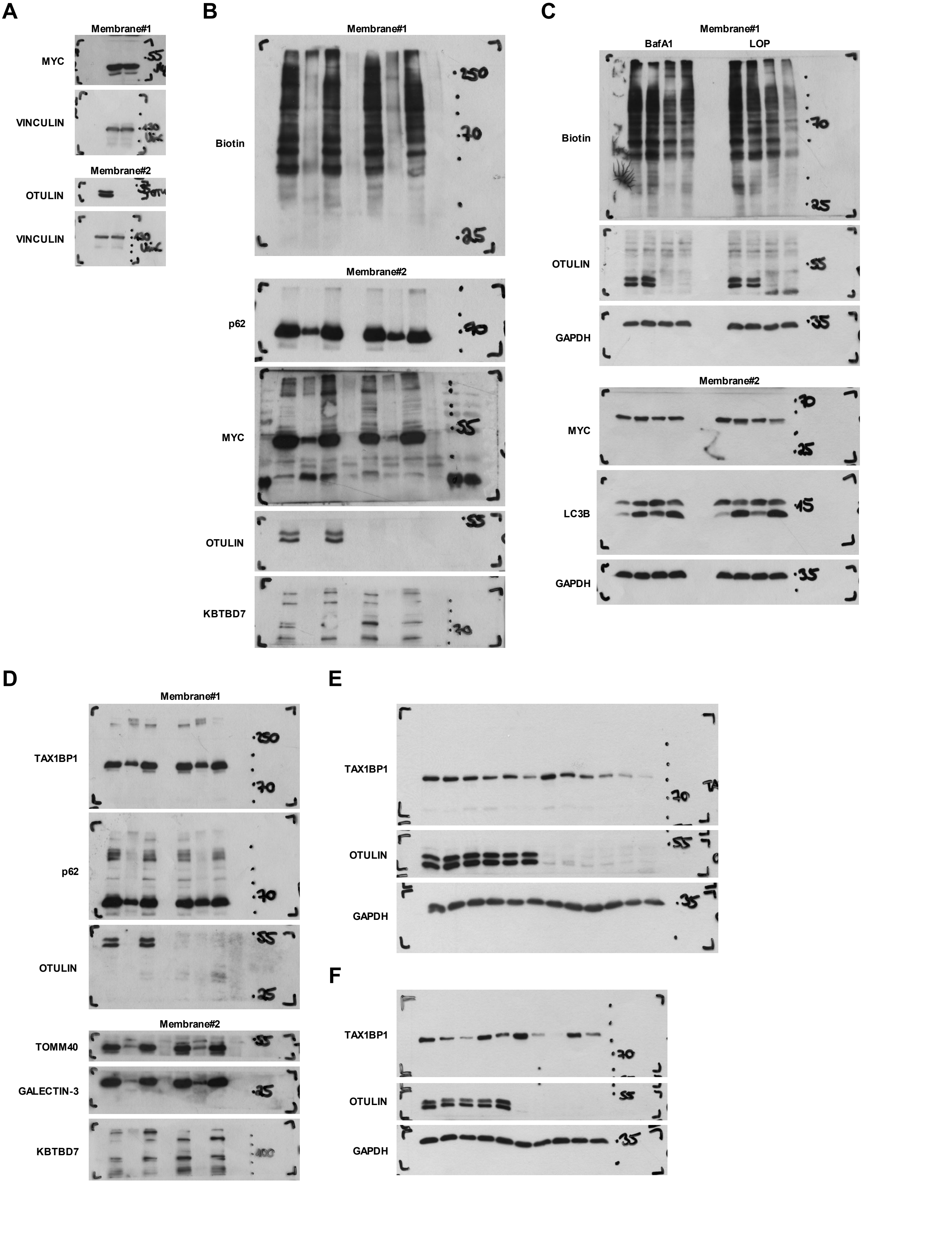


**Supplemental Figure 1**

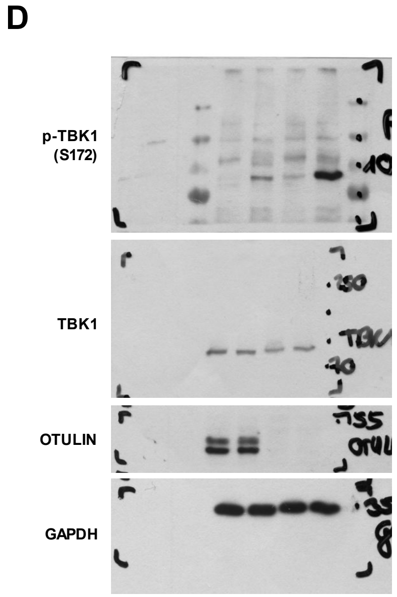


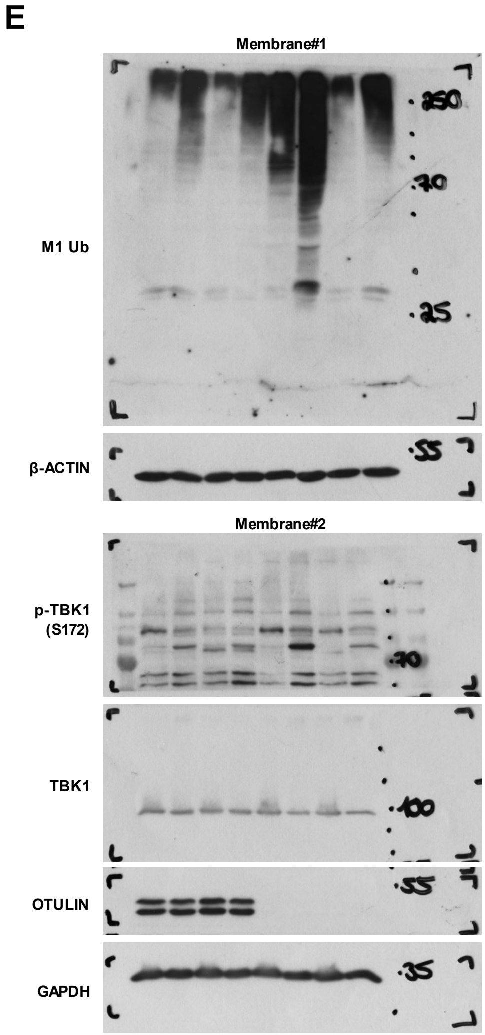


**Supplemental Figure 2**

### Supplementary Figures

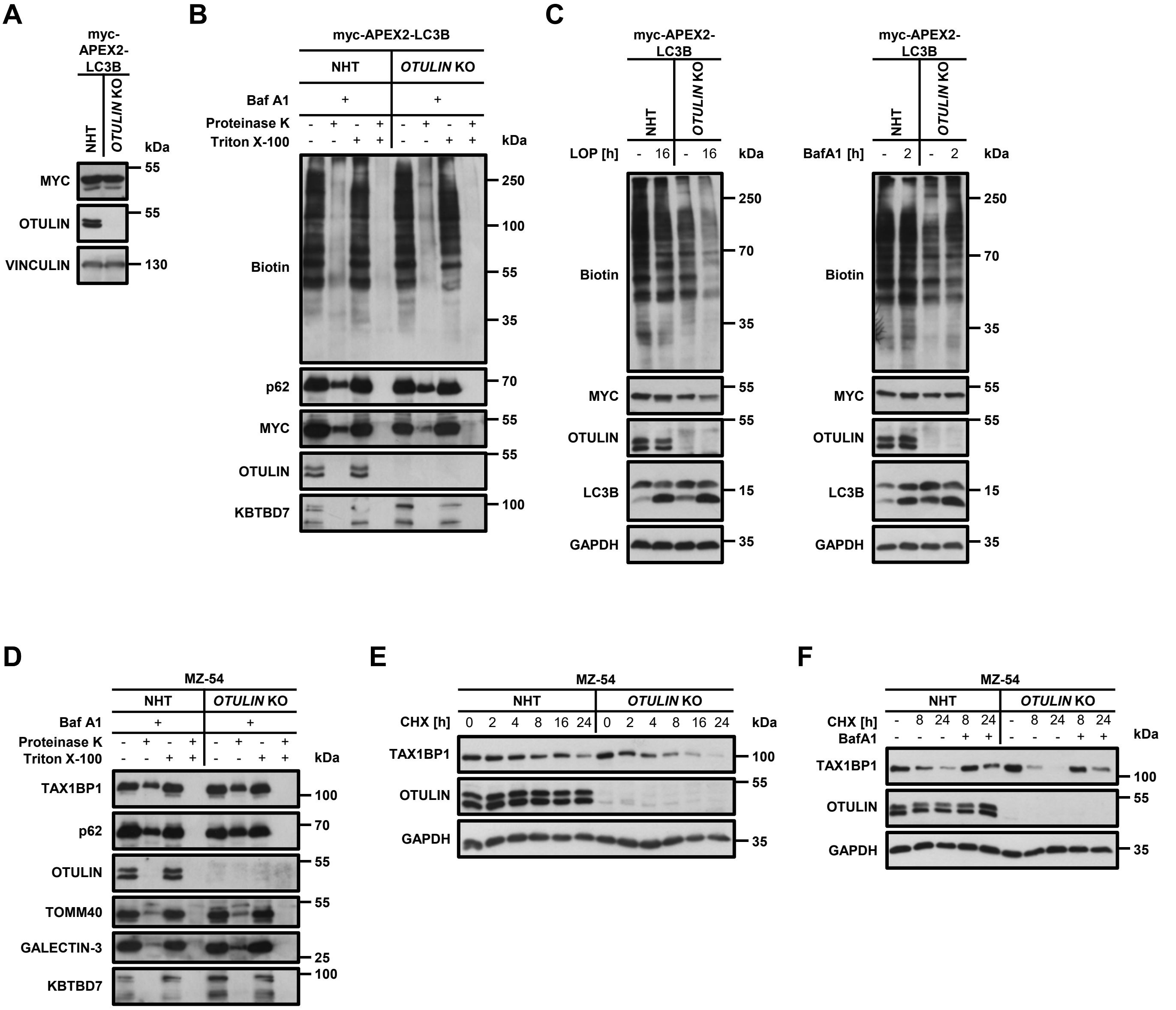


SF1 related to Figure 1

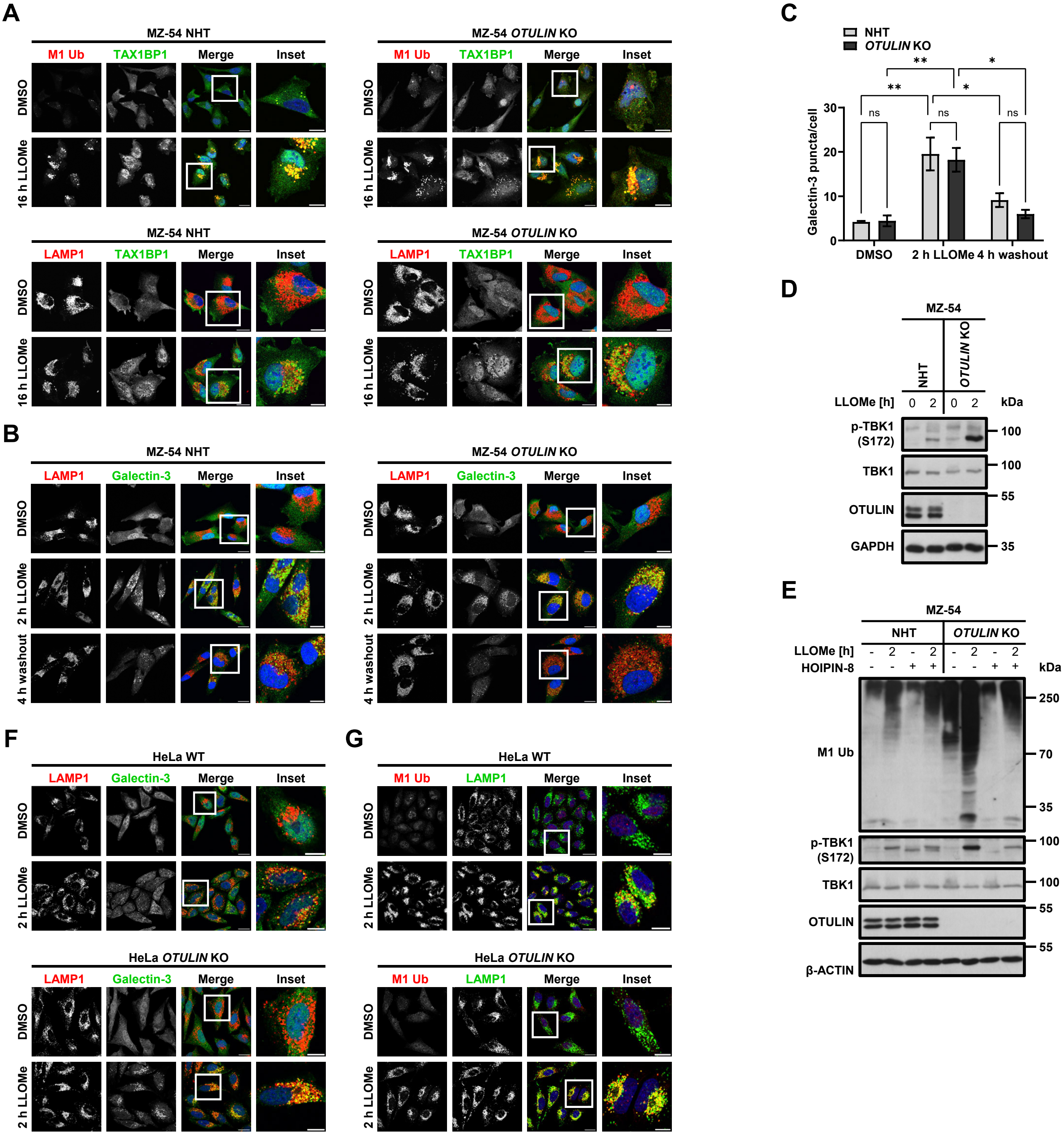


**SF2 related to Figure 2**

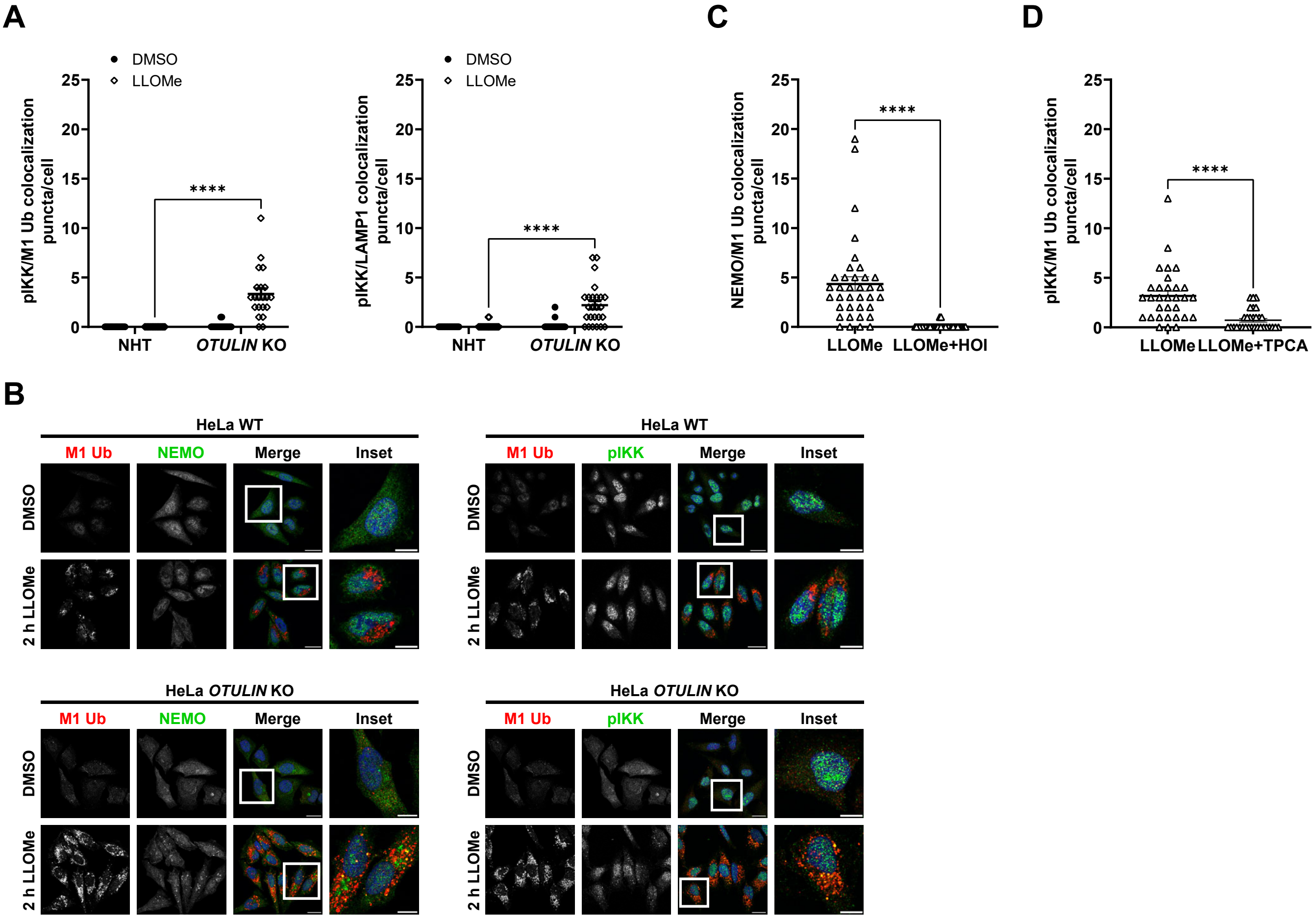


SF3 related to Figure 3

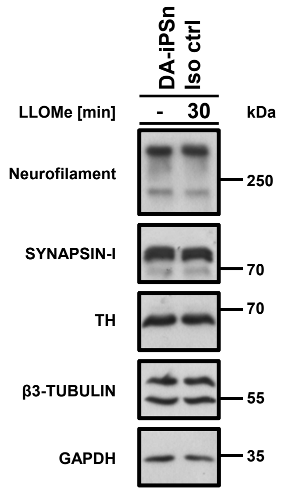


**SF4 related to Figure 6**
